# High-throughput protein characterization by complementation using DNA barcoded fragment libraries

**DOI:** 10.1101/2024.05.08.593210

**Authors:** Bradley W. Biggs, Morgan N. Price, Dexter Lai, Jasmine Escobedo, Luis Fortanel, Yolanda Y. Huang, Kyoungmin Kim, Valentine V. Trotter, Jennifer V. Kuehl, Lauren M. Lui, Romy Chakraborty, Adam M. Deutschbauer, Adam P. Arkin

## Abstract

Our ability to predict, control, or design biological function is fundamentally limited by poorly annotated gene function. This can be particularly challenging in non-model systems. Accordingly, there is motivation for new high-throughput methods for accurate functional annotation. Here, we use **co**mplementation of **aux**otrophs and DNA barcode **seq**uencing (Coaux-Seq) to enable high-throughput characterization of protein function. Fragment libraries from eleven genetically diverse bacteria were tested in twenty different auxotrophic strains of *Escherichia coli* to identify genes that complement missing biochemical activity. Although assay effectiveness ranged with respect to source genome, with 41% of expected enzymes recovered, even distant *E. coli* relatives like *Bacillus subtilis* and *Bacteroides thetaiotaomicron* showed success. Coaux-Seq provided the first experimental validation for 53 proteins, of which 11 are less than 40% identical to an experimentally characterized protein on an amino acid basis. Among unexpected function identified was a sulfate uptake transporter, an O-succinylhomoserine sulfhydrylase for methionine synthesis, and an aminotransferase. We also identified instances of cross-feeding wherein protein overexpression and nearby non-auxotrophic strains enabled growth. Altogether, Coaux-Seq’s utility is demonstrated, with future applications in ecology, health, and engineering.

## Introduction

Understanding the core metabolic functions of an organism is a critical step towards predicting its behavior and rationally manipulating function (Bordbar *et al*, 2014; Frioux *et al*, 2020; Widder *et al*, 2016). While a substantial amount of core metabolism is conserved, our ability to accurately annotate even well-known function is limited (Schnoes *et al*, 2009). For example, a recent study found that nearly one third of a diverse set of 127 bacteria were erroneously predicted to be auxotrophic for an amino acid (Price, 2023). Poor annotation at this level undermines approaches such as genome scale metabolic modeling, where gene annotation is foundational and errors introduced at the annotation step contribute significant uncertainty (D Ankrah *et al*, 2021; Bernstein *et al*, 2021). As various applications progress towards non-model organisms, the existence of isozymes, alternative biosynthetic pathways, and low amino acid identity homologs will continue to present a challenge. Failure to accurately identify core metabolic pathways will lead to misassignment of auxotrophies, misunderstanding of adaptive physiologies, and inferences of community dependencies that likely do not exist. Therefore, there is a need for new and high-throughput approaches to provide experimental evidence to accurately annotating these central pathways.

To this end, our laboratory has developed a suite of high-throughput functional genomics methods based on DNA barcoding and parallel fitness profiling, including transposon mutagenesis libraries (RB-TnSeq) (Price *et al*, 2018; Wetmore *et al*, 2015), dual-barcoded *E. coli* genomic fragment shotgun expression libraries (Dub-Seq) (Mutalik *et al*, 2019), and single-barcoded overexpression library screening in the anaerobe *Bacteroides thetaiotaomicron* (Boba-Seq) (Huang *et al*, 2022), along with development of CRISPRi tools for similar applications (Qi *et al*, 2013; Rishi *et al*, 2020). Here, we extend the DNA barcoding framework in functional genomics by employing shotgun expression libraries of diverse bacteria for the **co**mplementation of **aux**otrophs (Coaux-Seq, where biochemical function is “coaxed” from genetic material). By utilizing long-read sequencing to link genomic fragments to 20 nucleotide DNA-barcodes, we take advantage of facile and cost-effective barcode sequencing (BarSeq) to repeatedly assay genomic fragment libraries in different genetic contexts to test for the ability of contained gene material to encode protein(s) that complements a missing biochemical function in *E. coli*. Thus, we link genes from diverse bacterial genomes to known function. For this study, we generated 11 diverse bacterial genomic fragment libraries and tested these libraries in 20 different genetic knockout contexts. Beyond providing the first experimental evidence to validate the predicted biochemical function of 42 enzymes, we identified the function of 8 homologously divergent enzymes. Further, we identified unexpected function in 3 enzymes, including a sulfate uptake transporter from the TauE family, an O-succinylhomoserine sulfhydrylase for methionine synthesis (MetZ), and an aminotransferase.

## Results

To ascertain the genetic range of material capable of being successfully utilized in our proposed workflow, specifically in the context of *E. coli* expression, we sampled from genetically diverse bacteria. This included both gram-positive (e.g., *Bacillus subtilis*) and gram-negative (e.g., *Pseudomonas fluorescens*) strains, 3 different phyla (Bacillota, Pseudomonadota, and Bacteroidota), and 11 different genera. Classes represented include alphaproteobacteria (e.g., *Sphingomonas koreensis*), betaproteobacteria (e.g., *Xylophilus sp.*), gammaproteobacteria (e.g., *Lysobacter sp.*), bacteroidia (e.g., *Bacteriodes thetaiotaomicron*), sphingobacteriia (*Pedobacter* sp.), and bacilli (e.g., *B. subtilis*). The full set of 11 strains can be found in **Table S1** and in **Figure 1**, and summary information on the generated libraries can be found in **Table S1** and **Figures S1-11**. Several strains were obtained through recent isolations from the Oak Ridge Reservation, and their isolation conditions are described in detail in the methods. If not already available, the genome for each strain was sequenced and added to NCBI (full set of accession numbers in the methods).

**Figure 1.**
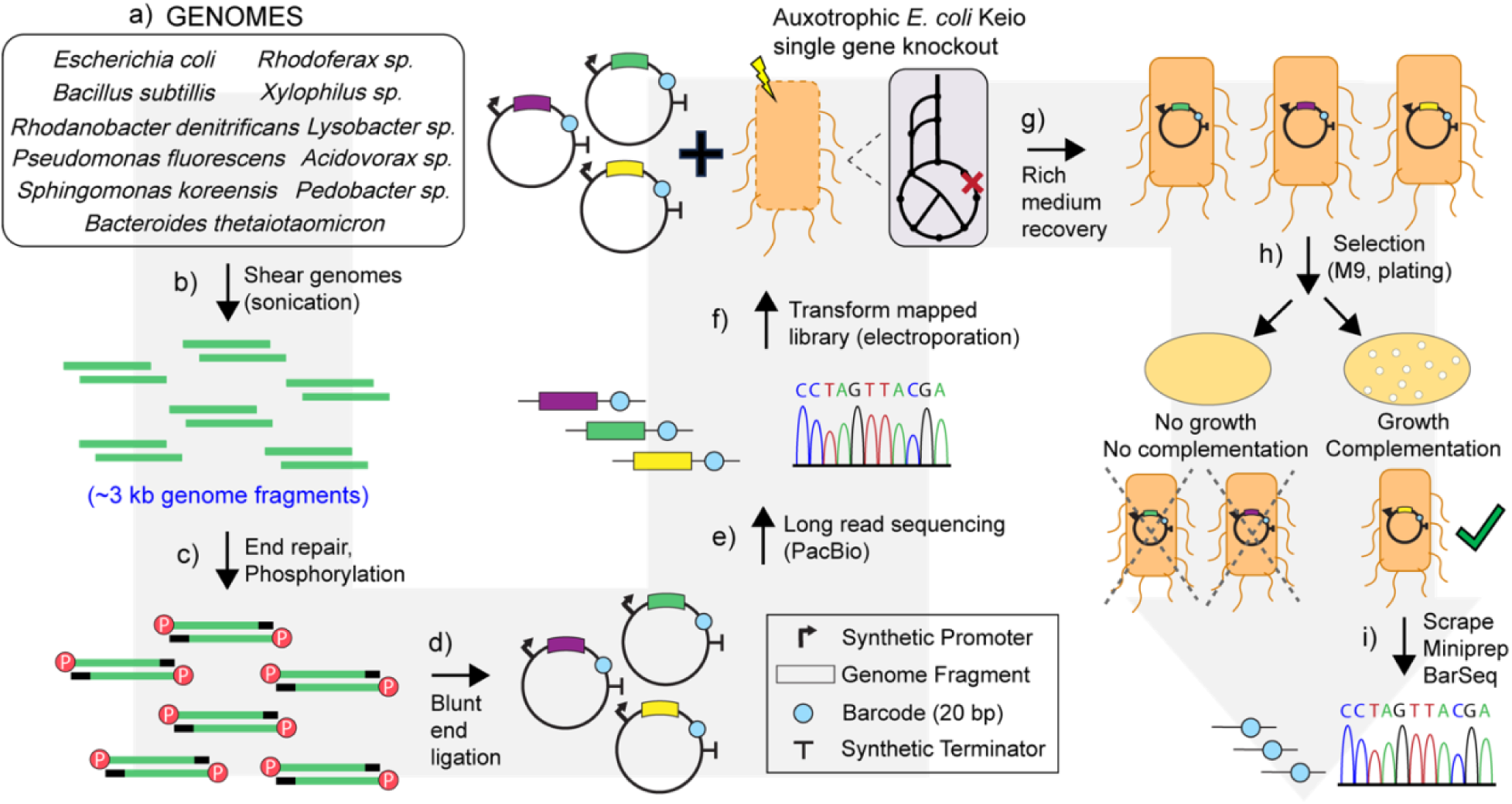
Coaux-Seq full workflow. Beginning in a) organisms are chosen and genomes extraction. In b) these genomes are sheared by sonication to ∼3 kb fragment size. In c) fragments are end repaired and phosphorylated. In d) blunt end ligation is used to ligate the repaired and phosphorylated fragments into the barcoded (random PCR generated 20 nucleotide segments) expression vector. In e) long read (PacBio) sequencing is used to map the fragments to the barcodes. In f) the libraries are transformed into a given auxotrophic strain. In g) transformed strains are recovered in rich medium (SOC, LB). In h) selection is conducted in minimal (M9, glucose) solid medium. In i) after selection, colonies are scraped, miniprepped, and BarSeq analysis is run.

The full experimental workflow for Coaux-Seq can be found in **Figure 1**. Briefly, extracted genomes are sheared by sonication to approximately 3 kb fragment size (**Figure S12** for example gel), end-repaired, phosphorylated, and blunt-end ligated into a barcoded expression vector. Barcoded expression vectors were generated by PCR using a primer with a degenerate overhang to introduce random 20 nucleotide-long sequences (barcodes) and to linearize the vector backbone prior to blunt-end ligation. For each of the 11 libraries, long read sequencing (PacBio) was used to link the ligated genome fragment to the unique barcode of its vector, following the library mapping workflow of Boba-Seq (Huang *et al*, 2022). By establishing this link between the fragment and barcode once, subsequent experiments only required more cost-effective barcode sequencing (BarSeq) to identify the genome fragment(s) contained. Mapped libraries could then be transformed, either individually or in groups, into different auxotrophic strains of *E. coli* and cultivated under selective conditions. For strains that grew under selection, the DNA could be extracted and sequenced by BarSeq.

Transformations were initially recovered in rich medium to determine transformation efficiency, both with respect to colony forming units (spotting assays) and with respect to the number of barcodes transformed (BarSeq). Rich medium recovered transformations were archived and could later be cultivated, pelleted, and washed (3x with 1x PBS) to remove nutrients before introduction to a selective condition. Selection was carried out in a defined minimal medium (M9) with 1% glucose as the sole carbon source. This medium also contained kanamycin for the genomic deletion selection marker (Keio collection (Baba *et al*, 2006)), chloramphenicol for the selection marker of the plasmid containing the gene fragment, and anhydrotetracycline (aTc) to induce expression of the Tet promoter. The Tet promoter was provided upstream of the genomic fragment insert to provide synthetic expression and was under regulation of the TetR repressor. The DNA barcode was added downstream of the fragment insert. A plasmid map has been provided with the supplemental materials. The vector used in this study contains a p15A origin of replication and an oriT origin of transfer to enable conjugation. **Figure S13** shows confirmation of inducible mRFP expression for this vector. The *E. coli* single gene knock-outs into which the libraries were transformed were auxotrophic, specifically unable to grow in M9 medium with glucose as a sole carbon source. Where growth was observed, colonies were scraped from the selective plates into sterile 1x PBS, pelleted, and plasmid miniprepped. Miniprepped plasmids were used as a template for BarSeq to determine which fragments were selected through this process.

Each of the 11 libraries were transformed individually into four different auxotrophic backgrounds (Δ*aroA*, Δ*proB*, Δ*metB*, Δ*thrB*) and as a combination of all 11 libraries into each of the 20 auxotrophic strains (Δ*aroA*, Δ*proB*, Δ*metB*, Δ*thrB*, Δ*cysA,* Δ*argG,* Δ*leuA,* Δ*trpA,* Δ*serA,* Δ*hisC,* Δ*metE,* Δ*ilvD,* Δ*pheA,* Δ*purE,* Δ*proA,* Δ*cysH,* Δ*hisG,* Δ*aroE,* Δ*pyrD,* Δ*ppc*) [Full information of auxotrophic strains in **Table S2**, full list of all library transformations in **Table S3**, and full list of all selections in **Table S4**]. This list of knock out strains was chosen based on the aforementioned criteria of being auxotrophic in minimal medium with glucose as a sole carbon source and to cover a range of chemistries within core metabolism. These activities included transferase (AroA; EC 2.5.1.19), kinase (ProB; EC 2.7.2.11), lyase (MetB: EC 2.5.1.48), synthetase (ArgG; EC 6.3.4.5), dehydratase (IlvD; EC 4.2.1.9), reductase/dehydrogenase (ProA; EC 1.2.1.41), carboxylase (Ppc; EC 4.1.1.31), and transporter (CysA; EC 7.3.2.2/5). The majority of knockouts came from amino acid biosynthesis (17), with two (purE, pyrD) from nucleotide biosynthesis and one (Ppc) from central metabolism (**Table S2**). All experiments were run at 37°C. Initial experiments were plated at multiple densities and optimized with respect to aTc induction level (**Figure S14**), with greater induction (transcription) tested to potentially compensate for weak heterologous ribosomal binding site (RBS) activity. From these experiments, 5x aTc (500 ng/mL) was found optimal (**Figure S14**), and subsequent experiments were run at both 1x (100 ng/mL) and 5x aTc, with 1x aTc used as a precaution for possible overexpression toxicity due to a 5x aTc induction level.

Initial library selections were performed on solid medium. This allowed for maintenance of a diversity of barcodes as measured by BarSeq and prevented consolidation to only a single or few barcodes, which was observed for liquid culture selections of libraries where the top two barcodes accounted for >99% of all reads in all three initial cases tested (**Tables S5-7**). Selection on agar plates, however, created a potential for false positives in the form of very small colonies that achieve minimal growth either due to carryover of nutrients even after multiple (3x) washing steps or from nutrients contained in agar by way of impurities. Upon scraping plates in preparation for plasmid miniprep and BarSeq, these faint colonies can contribute a small fraction of cell mass and thus DNA to subsequent experiments, necessitating cutoff criteria for enrichment to delineate successful fragments as defined below. These small faint colonies were observed even in control experiments run with red fluorescent protein (mRFP) in place of a genome fragment. However, false positive colonies did not grow beyond a small translucent state on solid medium. In addition, these false positive colonies did not grow in liquid medium (M9, 1% glucose with chloramphenicol, kanamycin, and aTc), save a single instance for Δ*pheA* as discussed further below.

### Total Library Hits

Using this workflow, we were able to identify complementation “hits,” meaning fragments containing genes for which the encoded biochemical activity improved the fitness of the auxotrophic *E. coli* in the selective condition (allowed growth). Fitness was defined as the log_2_ change in the relative abundance of a specific barcode in a given experiment. This was computed as the normalized log_2_ ratio of the number of reads for the barcode from the experimental sample (scraped up from an agar plate after growth in minimal medium) divided by the number of reads for that barcode from the control sample (used to inoculate the plate). As done previously(Huang *et al*, 2022), we defined high-confidence hits as fragments that provided significant benefit (fitness > 5 and z-like test statistic > 4) and either (1) an overlapping fragment provided a significant benefit in that same experiment or (2) the fragment provided a significant benefit in another experiment in the same mutant background (regardless of induction level).

Figure 2a shows an overview of the fragments with the potential benefits (fitness > 4) across all experiments. A blue “x” denotes a fragment that provided a significant benefit and the green diamonds indicate a fragment with high fitness that overlapped with another fragment with high fitness. Many of the inserts with high fitness overlap another insert with high fitness (52% of markers are the green diamonds). In addition, many of the inserts with high fitness show a benefit at both inducer concentrations (74% of markers that are above 5 for one axis are above 5 for both). With respect to overlapping fragments providing a fitness benefit, Figure 2b shows an example for TK06_RS12685 from *P. fluorescens* FW300-N2E2 in the context of Δ*hisC*. Highlighted in green are fragments that contain the whole TK06_RS12685 gene. Several fragments containing the whole gene show elevated fitness scores, while neighboring fragments that do not contain TK06_RS12685 do not show improved fitness scores.

**Figure 2.**
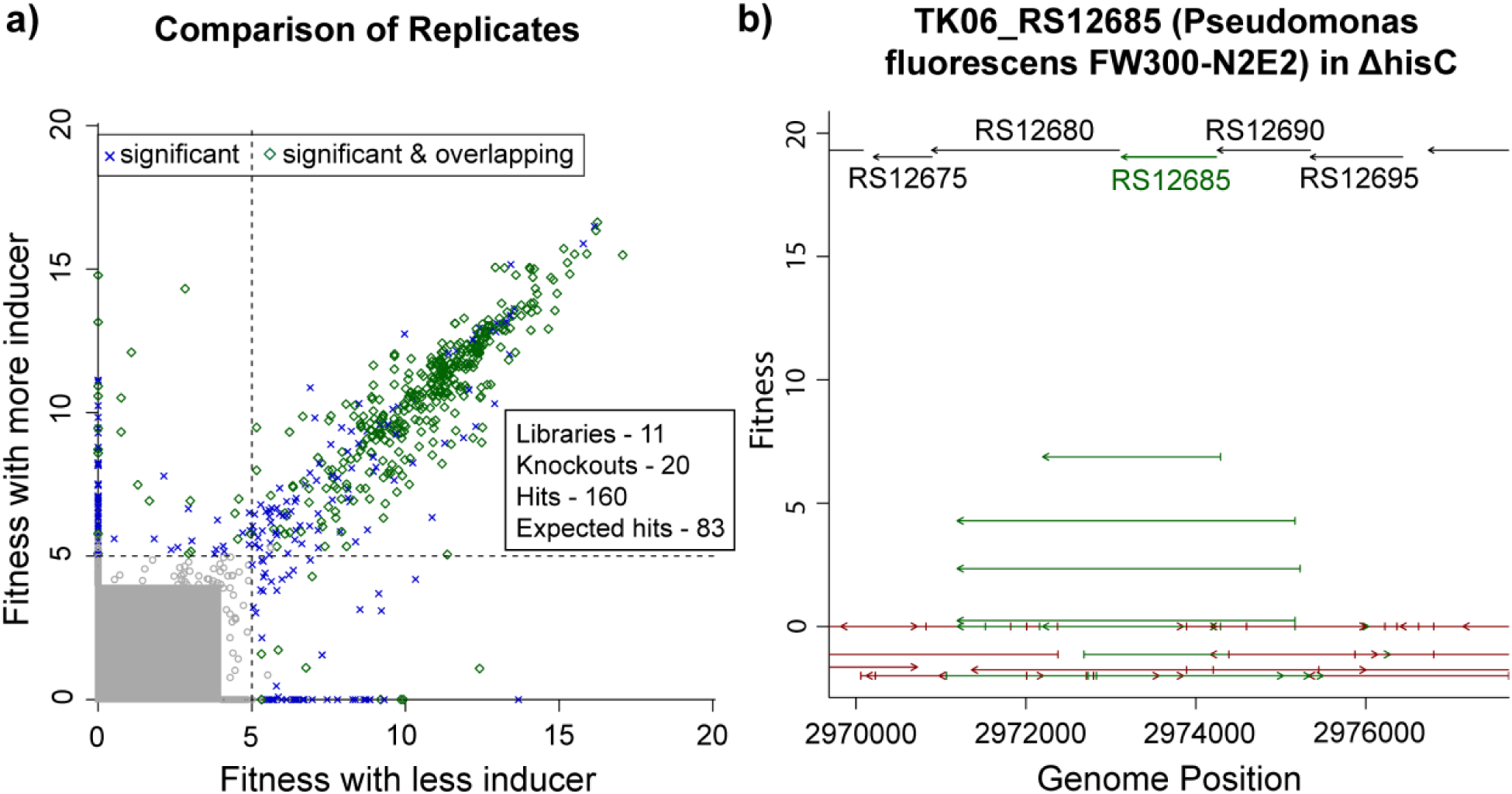
Coaux-Seq replicate comparison and an example case. In panel a), each point indicates the fitness values of a specific barcode from two experiments with the same mutant background and the same library (or mixture of libraries), but with different concentrations of the inducer. The fitness value from a lower concentration of inducer is on the x-axis. Points are highlighted if the barcode has a statistically-significant benefit in either experiment or if the barcode has a significant benefit and overlaps another barcode with a significant benefit. Points that are not statistically significant either fall within the gray box or are not highlighted (fitness values 4 to 5). Panel b) shows a case of a call of a hit for fragments associated with TK06_RS12685 from *P. fluorescens* FW300-N2E2 for activity in a Δ*hisC* auxotrophic background, where fragments that contain all of the gene (TK06_RS12685) are highlighted in green. Fitness values (y-axis) represent the average over two experiments. This specific example comes from an experiment where all 11 libraries were transformed together into the knockout background. If a fragment is higher on the y-axis, it showed a greater fitness improvement. The top of the plot shows a reference segment of the source genome. Length of the fragments below gives an indication of which genes were included in the given fragment. The arrow on the fragment indicates its directionality within the expression vector. For this case, all of the fragments with fitness above 0 are in the same orientation as the reference gene, indicating that they are all in the sense orientation with respect to the synthetic promoter.

In total, our experiments identified 838 instances of a fragment in the context a specific knockout background providing a significant fitness benefit in at least one experiment. Of these, 420 cases met the criteria of a high-confidence benefit. Combining the overlapping fragments among the 420 high-confidence hits, 160 regions were identified to a provide high-confidence fitness benefit. We associated these regions with genes by considering all genes that were contained within the insert with the highest fitness (averaging across experiments for that genetic background). Breaking down the 160 regions, 64 are associated with just one open reading frame (ORF), 75 are associated with two or more ORFs (47 with exactly two), and 21 do not contain an entire ORF. To identify causative genes, we first considered genes that lie within these regions and are expected to perform the missing function.

Although we did not perform exact replicate experiments, we repeated many of our experiments at more than one concentration of inducer, including experiments with the mixed library from 11 genomes for 19 of the 20 mutant backgrounds (because of a low number of colonies from the Δ*proA* transformation, the 1x and 5x aTc plates were combined). As mentioned, and demonstrated in Figure 2a, barcodes that had a benefit at one concentration usually had a benefit at the other concentration. This was almost always true if that barcode’s insert overlapped with the insert of another barcode that was significant in the same experiment (see green points in Figure 2a). In particular, if an insert had a significant benefit at the lower concentration, then it had a significant benefit at the higher concentration 88% of the time. For inserts whose benefit was confirmed by overlap, this proportion rose to 98%.

### Expected Hits

Based on the known genes of the source genomes and existing annotation tools, it is possible to create a list of enzymes that *a priori* would be expected to complement our auxotrophic knockout strains even if they have not been experimentally verified. To create a list of expected hits, we considered all of the knockouts except Δ*cysA*, as CysA is the ATPase subunit of a 4-component sulfate transporter complex and its association and functionality within the protein complex may present an independent issue beyond examining for a single enzyme catalytic function. From there, GapMind (Price *et al*, 2020), an annotation tool for amino acid biosynthesis that relies on experimentally characterized enzymes, was used to identify candidates for amino acid biosynthesis knockouts. For CysH and PyrD, TIGRFAMs (Haft *et al*, 2003) was used to identify candidates, using HHMer 3.3.1 and the trusted cutoff for the bit score (as provided by the TIGRFAMs curators). For PurE, we searched for homologs of the *E. coli* or *B. subtilis* PurE enzymes, which yielded a single candidate per genome. For Ppc, we only searched for homologs of *E. coli* phosphoenolpyruvate carboxylase, as *B. subtilis* does not have this enzyme. If a candidate enzyme was high-confidence (>40% amino acid identity to an experimentally validated enzyme that had the appropriate function, and where it was the only candidate in the genome), it was included as an expected hit.

There were some special cases that required additional effort, such as instances where there was more than one high-confidence candidate or if the original annotation was vague. For these, we used PaperBLAST (Price & Arkin, 2017), a tool that finds papers about homologs by using a combination of full-text search [EuropePMC (Gou *et al*, 2015)] and curated resources [Swiss-Prot (Bateman *et al*, 2023), BRENDA (Chang *et al*, 2021), EcoCyc (Caspi *et al*, 2020)], for confirmation. Steps without high-confidence candidates in the genome were manually examined, focusing on the lower confidence GapMind hits. This yielded two additional candidates. First, AAFF35_21465 from *Pedobacter sp.* FW305-3-2-15-E-R2A2 was identified as a likely AroA, as close homologs are essential proteins and it is 45% identical to HMPREF1058_RS13970 from *Phocaeicola vulgatus* CL09T03C04, which is cofit with chorismate synthase (AroC) in RB-TnSeq fitness data (Surya Tripathi, personal communication). Second, AAGF34_00495 of *Rhodoferax sp.* GW822-FHT02A01 was considered a likely SerA, as it is 68% identical to BPHYT_RS03150 from *Burkholderia phytofirmans* PsJN, which was identified as SerA using fitness data (Price *et al*, 2018). For MetB complementation, GapMind’s candidates for MetZ, which can substitute for both MetB and MetC, were also considered. However, these instances were excluded if the source organism was predicted to use O-acetylhomoserine in its methionine synthesis pathway instead of *E. coli’s* native O-succinylhomoserine pathway, as these candidates are unlikely to function appropriately in this background. For Ppc, two diverged homologs were included (LRK54_RS12075 of *Rhodanobacter denitrificans* FW104-10B01 and AAFF32_01485 of *Lysobacter sp.* FW306-1B-D06B). While these homologs are only 36-37% identical to characterized phosphoenolpyruvate carboxylases, they are likely to function as Ppc considering the functional residues are conserved, as identified using SitesBLAST (Price & Arkin, 2022). In addition, AAFF32_01485 from *Lysobacter sp*. FW306-1B-D06B is 58% identical to N515DRAFT_2010 from *Dyella japonica* UNC79MFTsu3.2, which is confirmed to be Ppc by its fitness pattern in RB-TnSeq data (Price *et al*, 2018). Overall, we identified 203 proteins expected to complement one of the 19 mutants, with 7-14 per mutant background and 14-22 per source genome.

### Expected Hit Recovery

Of the 203 expected complementing proteins, 80 were covered by a high confidence benefit fragment containing a full-length gene, and 3 were covered by a nearly full-length gene (41% recovery of expected hits), representing 52% of all of the high-confidence hits. Perhaps as would be anticipated, as the contained expression components (promoters and RBSs) of the *E. coli* fragments are likely to work well in the *E. coli* knockout host, the recovery rate of expected hits was highest for *E. coli* (95%). *P. fluorescens* FW300-N2E2 showed the second highest recovery rate (68%). All other libraries showed a lower recovery rate.

In an effort to understand why some expected hits were not recovered, we looked into several possible explanatory effects. First, we checked to see if a fragment containing the expected hit was present in the generated libraries, and a full-length gene was found in the libraries for 75% of the expected hits. Second, we examined whether the expected gene was contained in the library at t_0_, to determine if the fragment transformed successfully. Of the remaining 152 genes covered by a fragment, only 132 of them were seen at t_0_. Moreover, as would be expected, inserts that contain a should-be beneficial protein but were not detected in the t_0_ samples (0 reads for their barcode) are much less likely to show a benefit compared to inserts that are detected (2% vs. 50%), with the 2% likely representing fragments that were transformed at low abundance and missed in the t_0_ sequencing. Together these effects accounted for roughly 70 of the 120 not recovered hits.

Next, we examined whether relatedness to *E. coli* was explanatory for success. This analysis considered fragment recovery percentage for fragments detected at t_0_ and in the sense orientation with respect to the synthetic promoter. We found no clear correlation between fragment recovery success and fragment genome origin relatedness to *E. coli*. For example, one of the most distantly related organisms, *B. subtilis,* showed good recovery of expected complementing genes (**Figure S15,** 72%). At the same time, the gammaproteobacteria *Lysobacter sp.* FW306-1B-D06B and *R. denitrificans* FW104-10B01 showed poor recovery (35% each), while other gammaproteobacteria (*E. coli* and *P. fluorescens* FW300-N2E2) were among the best with respect to recovery (87% and 78%, respectively). Similarly, the betaproteobacteria *Rhodoferax sp.* GW822-FHT02A01 and *Xylophilus sp.* GW821-FHT01B05 showed poor recovery (40% and 36%), but other betaproteobacteria, including *Acidovorax sp.* FHTAMBA, showed good recovery (68%). While the alphaproteobacteria *S. koreensis* showed poor recovery of expected hits (35%), it was the only representative. All the other genome libraries besides *E. coli* showed between 60-70% percent recovery.

Next, we considered possible expression effects. Among inserts that were detected in the t_0_ samples and are expected to provide a benefit, those that have the gene in the same orientation as the synthetic promoter are nearly 2x more likely to demonstrate a benefit (66% vs. 35%). Following, we investigated expression impacts of ribosomal binding sites (RBS). Because of the manner of assembly of the libraries (random shearing), our expression vector contained a synthetic promoter but not a synthetic RBS, as the gene could be located anywhere in the approximately 3 kb region of the insert fragment and in either orientation. Therefore, expression of fragment contained genes depends on native RBS function. We used the OSTIR calculator (Roots *et al*, 2021) to determine expected expression strength of source genetic material RBSs. The only library with significantly weaker RBS strength on average was *S. koreensis* (mean RBS strength of 1.3, as compared to 2.4 across other libraries; P = 0.007, t-test with Bonferroni correction for 11 libraries tested), and this library did perform poorly (35% recovery). However, *S. koreensis* contains a relatively high number of leaderless transcripts (Lomsadze *et al*, 2018), which would inherently deliver a low RBS score, as no RBS would be present, and are unlikely to be expressed in *E. coli* anyway. When considering only inserts that cover the entire gene, were oriented correctly with respect to the synthetic promoter, and were detected in the t_0_, the mean RBS strength was 2.36 for inserts with significant benefits and 2.37 for those without (P = 0.89, t-test). Overall, the RBS strength prediction did not correlate with individual protein complementation success. Lastly, we looked at genome GC content. Several of the genomes with low success rates have much higher GC content than *E. coli*, which could lead to poor expression, possibly via codon usage. However, comparing percentage of rare codons or codon adaptation index [CAI (Sharp & Li, 1987)] for genes that did or did not complement showed no clear differences.

Considering the 83 expected hits recovered, all 11 libraries were represented with at least two hits each, with the distribution favoring *E. coli*, *P. fluorescens* FW300-N2E2, and *B. subtilis* (Figure 3a). And while RBS strength was not predictive of an individual protein’s success rate to complement, the RBS strength calculations did show that *B. subtilis* had the highest average score (3.6), which may be due to its low GC content and could contribute to the somewhat unexpected strong performance in *E. coli*. For comparison, *E. coli* had the second highest average score (3.0). Lastly, as expected, a majority of the hits were in the sense orientation with respect to the synthetic promoter (Figure 3b, 65%). Excluding the *E. coli* gene fragments, for which natively contained promoters would be expected to perform well in our assays, 67% of hits were in the sense orientation with respect to the synthetic promoter.

**Figure 3.**
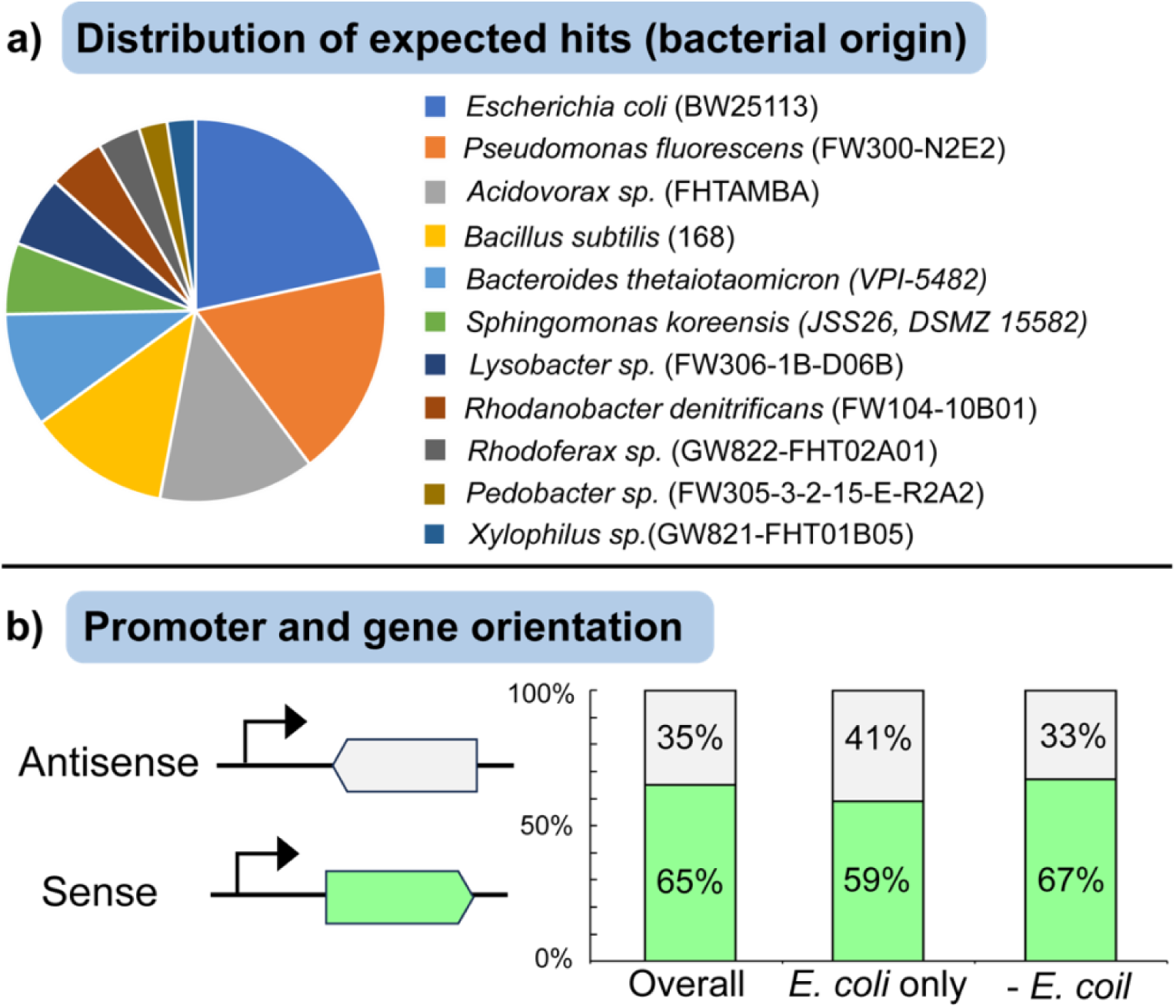
Number of expected hits recovered from each library and the orientation of inserts that have a statistically significant benefit and contain a gene expected to confer a benefit in that experiment, without considering confirmation by replicates or other inserts. Panel a) shows the breakdown of the genome of origin for the recovered expected hits. Panel b) shows the breakdown of the orientation of the complementing gene with respect to the synthetic promoter among inserts that have a statistically significant benefit and contain a gene expected to confer a benefit in that experiment.

### Diverged hits

Additionally, we categorized a set of “diverged” hits (10 in total) that could have been expected to complete the missing function, but with caveats to high-confidence assignment. Hits were considered “diverged” if they were similar to experimentally-characterized proteins that have the missing activity (and not more similar to proteins known to have other functions instead), but are <40% identical on an amino acid basis to any characterized protein with that activity in curated databases (Swiss-Prot (Bateman *et al*, 2023), MetaCyc (Caspi *et al*, 2020), BRENDA (Chang *et al*, 2021), and fitness browser reannotations (Price *et al*, 2018)). **Table S8** provides a full list of such cases. For example, AAGF34_01100 from *Rhodoferax sp.* GW822-FHT02A01 successfully complements *E. coli* Δ*hisC*, while only being 31% identical to hisC Q8R5Q4 of *Thermoanaerobacter tengcongensis*. At the same time, GW822FHT02A01_127 is found in a histidine biosynthesis operon, so this behavior could be expected. As another example, *B. subtilis* MetI (MetB-like) was found to complement Δ*metB*, even though *B. subtilis* uses acetylated intermediates and *E. coli* uses succinylated intermediates. This complementation was previously reported to work, even though the enzyme has no detectable activity on O-succinylhomoserine (Auger *et al*, 2002).

### Identifying novel enzymes and transporters

With respect to the remaining 75 hits that were not associated with an *a priori* expected complementation of the auxotrophic activity or our diverged set, we chose to follow up with 14 cases for validation (**Table S9**). Fitness plots for the associated fragments can be found in **Figures S16-29** and Figure 2b. For each of these cases, the identified gene fragment was cloned from the host genome into the expression vector exactly as it was found in the library fragment. Following, it was tested directly in the relevant auxotrophic *E. coli* host, selected on solid medium, inoculated into liquid medium, the plasmid was miniprepped, and sent for Sanger sequencing to confirm complementation and avoid false positives.

Of these 14 cases tested, 6 were found not to complement, 4 were found to benefit by cross feeding, and 4 were found to successfully complement. **Figure S30** shows a breakdown of the fitness scores and frequency of overlap for each of these categories along with the expected and diverged hits, and shows that the 6 cases found not to complement trend towards lower fitness scores and lack of overlapping fragments. Cross feeding and successful cases are described in further detail below. Among the cases that were confirmed to not complement were the putative transporter BSU_32380 of *B. subtilis* in the context of Δ*cysH*, BW25113_RS20525 (*ppc*) of *E. coli* in the context of Δ*proB*, TK06_RS22845 (*ilvD*) of *P. fluorescens* FW300-N2E2 in the context of Δ*proB*, and TK06_RS26365 of *P. fluorescens* FW300-N2E2 in the context of Δ*pyrD*. Likewise, the two putative major facilitator superfamily (MFS) transporters LRK54_RS17455 of *R. denitrificans* FW104-10B01 and TK06_RS20405 of *P. fluorescens* FW300-N2E2 were both found to not complement in the context of Δ*pheA*.

Several additional unexpected hits were found not to complement when individually expressed in the appropriate auxotrophic strain, but among a subset of these we recognized the possibility of a fitness benefit for the contained fragment in the context of a nearby non-auxotrophic strain cross feeding a needed metabolite. For example, the unsuccessful fragments tested in the context of Δ*ppc* were putative dicarboxylate transporters or symporters. When spotted on a plate proximal to wild-type *E. coli*, strains overexpressing these transporters showed improvement of growth compared to a control (mRFP in the place of a gene fragment) (**Figure S31**). These results indicate that the overexpression of these transporters provides a growth benefit, which occurs likely through improved uptake of dicarboxylic acids. Of possible candidate metabolites, succinate is most likely, as *E. coli* is known to secrete it (Clark, 1989) and it provides an alternate entry point to the TCA cycle. Similarly, overexpression of BT_RS23500 introduced to Δ*trpA* showed a growth benefit in cross feeding assay, contrasting a negative control (mRFP), and only in the case of spotting with a wild-type *E. coli* (**Figure S32**). BT_RS23500 is 59% identical to TrpB2 from *Thermotoga sp.* (Q9WZ09), an enzyme that forms tryptophan from indole (Hettwer & Sterner, 2002). This suggests that indole secreted from the wild-type *E. coli* (Wang *et al*, 2001) could be taken up by the auxotrophic strain and converted to tryptophan by BT_RS23500 to alleviate the loss of TrpA’s role of cleaving indole-3-glycerol phosphate (IGP) into glyceraldehyde 3-phosphate (GAP) and indole as part of tryptophan biosynthesis. It is somewhat surprising that *E. coli’s* native TrpB cannot provide this same benefit, but BT_RS23500 may possess superior kinetics. Its homolog, *Thermotoga* sp. trpB2 (Hettwer & Sterner, 2002) has a K_M_ for indole below 1 μM, while TrpB from *E. coli* (in the absence of TrpA) has a K_M_ of 14 μM. Together, these results indicate the importance of considering cross feeding in analyzing complementation assay results.

The two MFS transporters identified in the context of Δ*pheA* (LRK54_RS17455 of *R. denitrificans* FW104-10B01 and TK06_RS20405 of *P. fluorescens* FW300-N2E2) were also tested for cross feeding growth benefit. Neither demonstrated a clear cross feeding benefit, though strains overexpressing these transporters showed greater plate-based growth than the negative control (**Figure S33**). Unexpectedly, negative controls in these assays sometimes grew to a significant amount (**Figure S33**, agar plate on the right in the figure as an example). As previously mentioned, in other knockout contexts, negative controls would sometimes show small faint colonies on solid medium, but did not grow in liquid culture. However, if given extended periods of time (∼1 week), negative controls in Δ*pheA* grew in liquid culture. The long-lag time growth was different compared to strains overexpressing truly complementing fragments like TK06_RS12685 of *P. fluorescens* FW300-N2E2 and BT_RS19865 of *B. thetaiotaomicron*, which grew in 1-2 days. These observations led us to examine this behavior further.

To understand what might be occurring, we sequenced the genome of a Δ*pheA* negative control strain that grew to look for a compensating mutation that may facilitate this growth. However, we found no clear candidate mutations that might provide a benefit (**Supplemental File** for all differences between wild-type *E. coli* BW25113 and this strain). PheA is a bifunctional enzyme with both chorismate mutase and prephenate dehydratase activity. Because *E. coli* possesses alternative enzymes for the chorismate mutase activity, Δ*pheA* complementing fragments are generally expected to be carrying out the prephenate dehydratase activity. Interestingly, this activity can occur spontaneously, particularly at low pH (Cerutti & Guroff, 1965; Kishore *et al*, 1999). Therefore, it is possible that over extended periods of time, non-enzymatically catalyzed reactions are sufficient to provide growth for negative controls. As we consistently allowed multiday windows for growth in confirmation experiments, we observed Δ*pheA* growth when others may not have. These results indicate that spontaneous or even slow alternative enzymatic reactions should be taken into account as possible causes for cell growth when analyzing complementation assay data, particularly when extended periods for growth are allowed (such as for follow up experiments). Nevertheless, our complementation assays successfully identified 8 of the expected 12 PheA proteins (including *E. coli* PheA) as complementing this mutant, so this phenomenon did not entirely undermine the assay.

The four remaining unexpected hits were confirmed to complement. Among the confirmed complementing fragments, two examples came from the context of Δ*cysA*. Sulfur is an essential element for microbial metabolism, and microorganisms, including *E. coli*, utilize sulfate importers to obtain it. *E. coli* employs a sulfate/tungstate uptake transporter (SulT) family complex CysUWA-CysP/sbp, where CysP and sbp are alternate periplasmic substrate-binding proteins (Figure 4) (Aguilar-Barajas *et al*, 2011). Within this ABC importer complex, CysA is responsible for ATP energy coupling (Aguilar-Barajas *et al*, 2011). Accordingly, when testing fragments for complementation in Δ*cysA*, one would expect to identify hits that are CysA homologs. Beyond identifying *E. coli* CysA to complement itself, we only found two other fragments that allowed for *E. coli* growth in the context of Δ*cysA*, neither of which contained a CysA homolog. First, and perhaps as could be expected, we found that CysP from *Bacillus subtilis* replaces the missing functionality to allow for *E. coli* growth. CysP, a sulfate permease from the phosphate inorganic transporter (PiT) family (Mansilla & De Mendoza, 2000), complements the lost activity of CysA by replacing the entire CysUWTP complex function. More interestingly, a previously unannotated transporter, the TauE homolog TK06_RS10770 of *P. fluorescens* FW300-N2E2, was also confirmed to restore *E. coli* Δ*cysA* growth. The TauE family includes a sulfite exporter and a sulfoacetate exporter, but to date has not previously been linked to sulfate uptake. That stated, TK06_RS10770 is 49% homologous to Ac3H11_578 from *Acidovorax sp*. GW101-3H11, which is an operon with sulfate assimilation genes. Moreover, data from transposon mutants also links Ac3H11_578 to sulfate assimilation. Across 140 RB-TnSeq experiments, the fitness pattern of Ac3H11_578 is most correlated with a subunit of sulfate adenylyltransferase (linear correlation = 0.94) (Price *et al*, 2018). Combining the complementation data, RB-TnSeq data, and genome context, we can conclude that TK06_RS10770 and Ac3H11_578 are sulfate uptake transporters.

**Figure 4.**
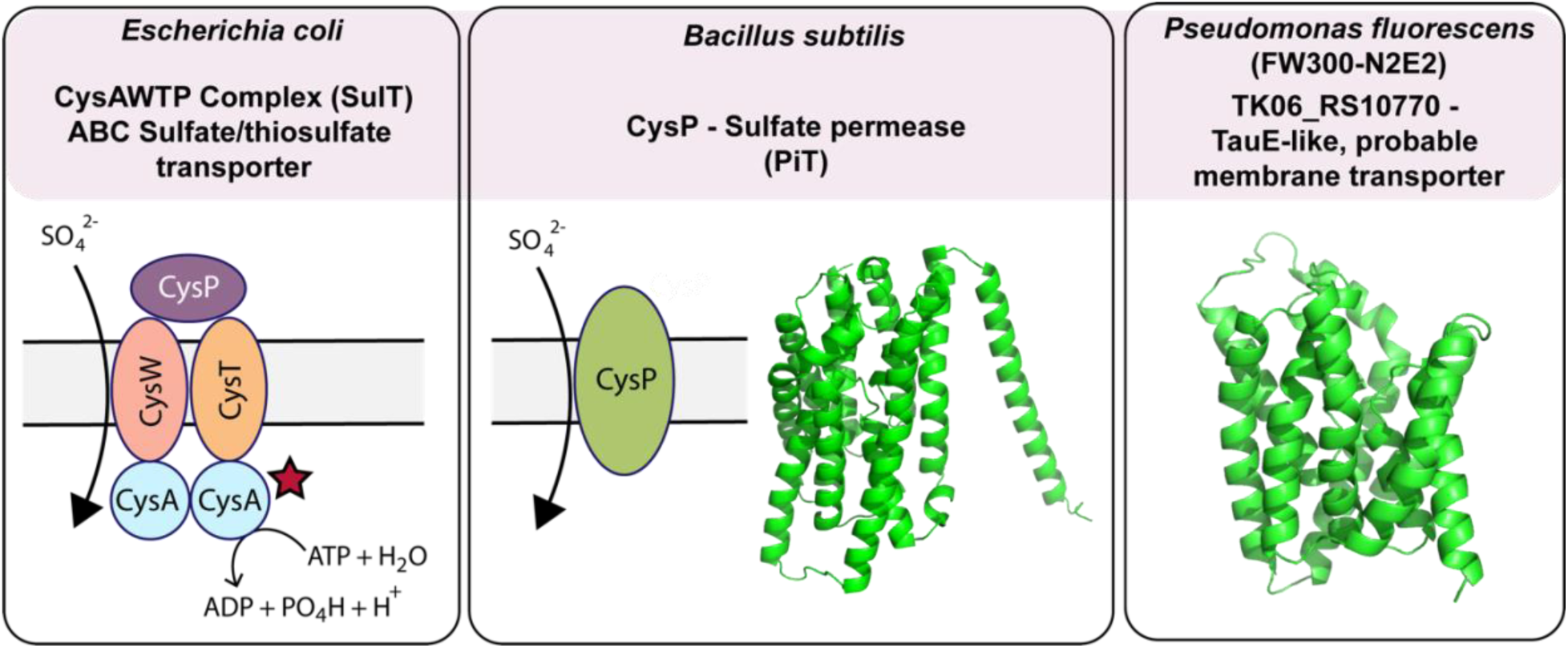
Complementation of CysA activity by *Bacillus subtilis* CysP and by novel sulfate transporter TK06_RS10770 from *Pseudomonas fluorescens* FW300-N2E2. First panel shows the native *E. coli* system. Second panel shows the function of the sulfate permease (CysP) from *B. subtilis*, along with its AlphaFold predicted structure. The final panel shows the AlphaFold predicted structure for TK06_RS10770 of *P. fluorescens* FW300-N2E2.

A second example of a confirmed hit is that of LRK54_RS05660 of *R. denitrificans* FW104-10B01 found in the context of Δ*metB*. MetB is a cystathionine gamma-synthase that, along with cystathionine beta-lyase (MetC), is involved in the essential two-step process of forming L-homocysteine from L-cysteine and O-succinyl-L-homoserine as part of *E. coli’s* methionine synthesis pathway (Figure 5). By homology, LRK54_RS05660 is related to both *E. coli* MetB (41%) and *E. coli* MetC (28%), with its AlphaFold structure resembling MetB (Figure 5). Yet, its closest biochemically-characterized homolog is cystathionine gamma-lyase from *Pseudomonas aeruginosa* (65%) (Pedretti *et al*, 2024). Moreover, RB-TnSeq data for LRK54_RS05660 does not show auxotrophic phenotypes (Adam Deutschbauer, personal communication), leaving unclear its function. As some microbes utilize an alternative pathway from O-succinyl-L-homoserine to L-homocysteine by way a single step with MetZ (Figure 5), we tested to see if LRK54_RS05660 were able to also complement Δ*metC,* another Keio collection knockout, in addition to Δ*metB*, which might indicate it was instead a MetZ. LRK54_RS17455 was not found able to complement Δ*metC*.

**Figure 5.**
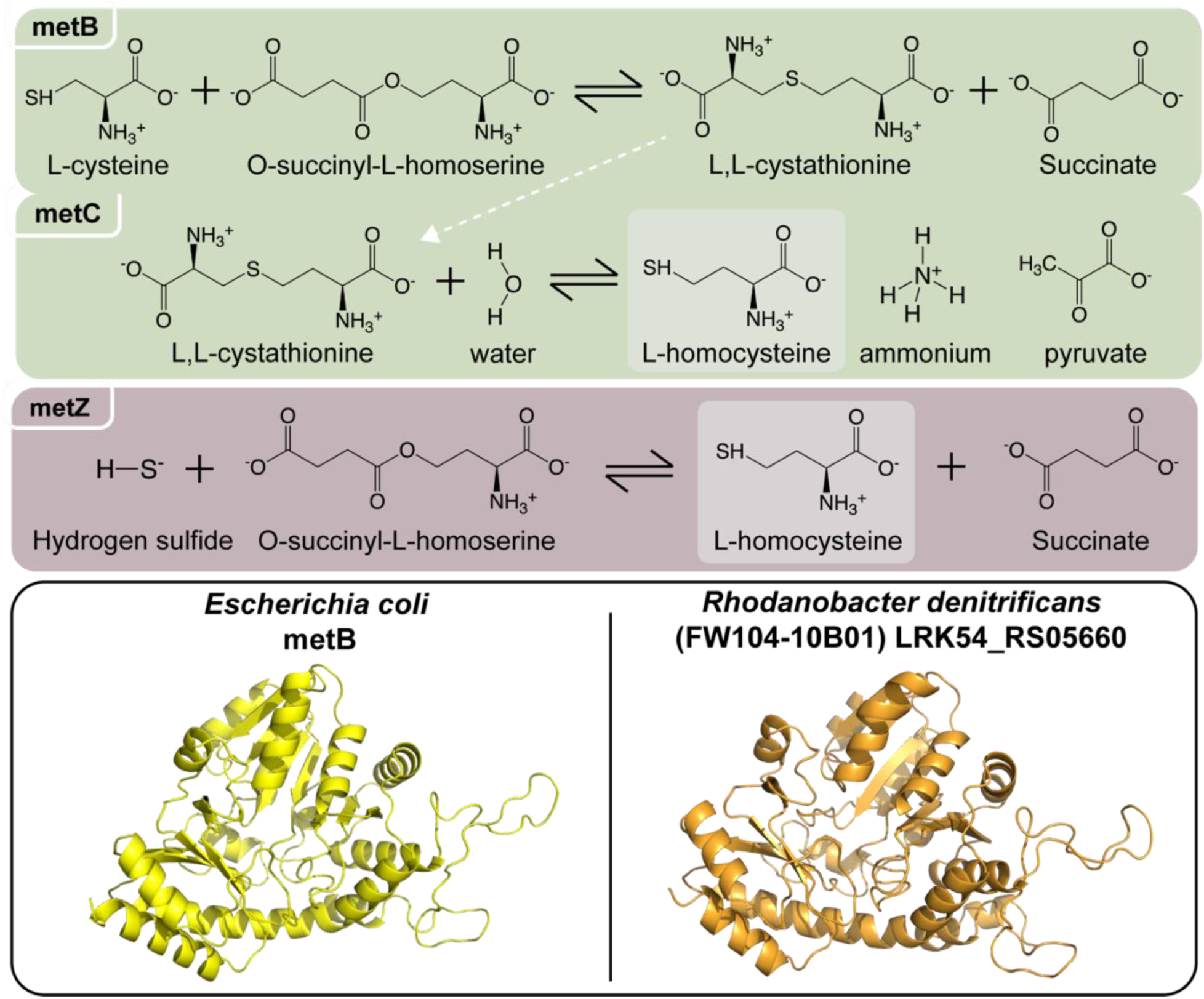
Identification of a protein, LRK54_RS05660 of *R. denitrificans* FW104-10B01 that complements Δ*metB* knockout. The top of the figure shows the two dominant pathways from homoserine to homocysteine used by bacteria, the two step (MetBC) and one step (MetZ) pathways. The lower portion of the figure shows crystal structure of *E. coli* MetB and the AlphaFold predicted structure for LRK54_RS05660, which bears structural similarity.

In contrast, another gene that complements Δ*metB,* AAFF19_12795 of *Acidovorax sp.* FHTAMBA, is predicted as either a methionine gamma-lyase or MetB, and we experimentally found that AAFF19_12795 was able to fill the role of a MetZ. AAFF19_12795is 90% identical to Ac3H11_2452 from *Acidovorax sp.* GW101-3H11, which is strongly cofit with homoserine O-succinyltransferase [*metA*, r = 0.98 (Price *et al*, 2018)]. Interestingly, neither FHTAMBA nor GW101-3H11 has a strong candidate for MetC, and the best candidate in the GW101-3H11 strain (Ac3H11_34) is more similar to methionine gamma-lyases and is not important for growth in minimal media (all RB-TnSeq fitness values > 0) (Price *et al*, 2018). Because of the lack of a clear MetC enzyme in these genomes, we hypothesized that both AAFF19_12795 and Ac3H11_2452 were MetZ (Foglino *et al*, 1995). Our initial experimental evidence of AAFF19_12795 complementing Δ*metB* leaves possible either MetB or MetZ functionality. Therefore, we tested AAFF19_12795 for Δ*metC* complementation, and found that it indeed complemented Δ*metC*, providing evidence that it is likely MetZ. Though this general activity could have been expected because of homology, additional assays were necessary to clarify its protein function. These results indicate the importance of considering possible alternative functions when analyzing complementation assay data.

As a final example, TK06_12685 of *P. fluorescens* FW300-N2E2 was confirmed to complement Δ*hisC* (Figure 6). TK06_RS12685 exists within a conserved operon containing aromatic amino acid biosynthesis genes for both tyrosine and phenylalanine synthesis, along with SerC and cytidylate kinase, suggesting a possible role related to amino acid biosynthesis. It is 81% identical to PA3165 (HisC2) of *P. aeruginosa*. And while HisC2 is not required for histidine synthesis (Wang *et al*, 2020), TK06_12685 also has 43% homology to *B. subtilis* HisC, providing further possible connection to this role. Interestingly, TK06_RS12685 also has 55% homology to BPHYT_RS14905 of *Burkholderia phytofirmans* PsJN, which is thought to be a phenylalanine transaminase because it is important for fitness during growth on phenylalanine (Price *et al*, 2018). Together, this evidence might suggest that, based on sequence similarity, TK06_RS12685 is a transaminase for phenylalanine or histidinol phosphate. Our data strongly suggests that it can use histidinol phosphate as a substrate, even though this might not be its physiological role. In summary, these three examples show that beyond validating expected function, Coaux-Seq successfully uncovered novel biochemical function.

**Figure 6.**
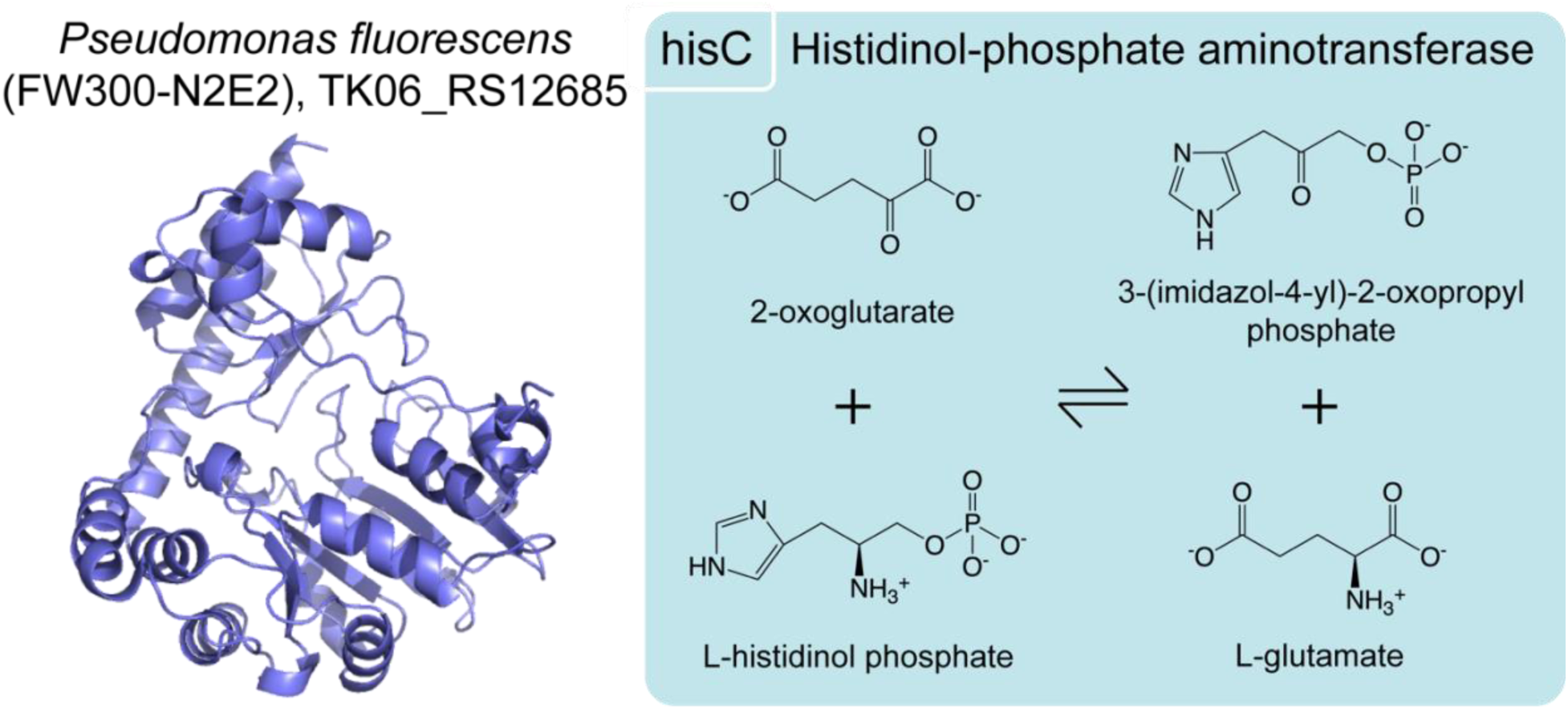
HisC activity from TK06_RS12685 of *Pseudomonas fluorescens* FW300-N2E2. One the left is the AlphaFold predicted structure of TK06_RS12685. On the right is the activity of HisC from *E. coli* that it is being complemented.

### Results summary, classifications

Summarizing the proteins identified across the different categories, we provided first experimental evidence for function of 53 proteins, of which 42 would have been reasonably expected to perform the identified function owing to their at least 40% amino acid homology to a protein with experimentally verified function. To assign this classification, we assumed that all *E. coli* and *B. subtilis* proteins are already characterized, although some do not exist in curated databases (e.g. ThrB from *B. subtilis*). Similarly, we excluded proteins that we previously identified by way of mutant phenotypes (Price *et al*, 2018), of which there were 6. Lastly, we compared to PaperBLAST’s database of characterized proteins to ensure that there were no other characterized proteins. This analysis identified two that were very close to a previously characterized protein (AAFF19_12795 is 92% identical to Ac3H11_2452, which is the MetZ discussed in the text, and TK06_RS20265 is 92% identical to Q3KK58 of *P. aeruginosa*, which is characterized as TrpA). Together, the complementation data contributed to 5 diverged enzymes being added to GapMind (Price *et al*, 2020) (BT2186/BT_RS11065 of *B. thetaiotaomicron* VPI-5482 as AroA, AAFF19_05770 of *Acidovorax sp*. FHTAMBA as HisC, AAFF35_11135 of *Pedobacter sp*. FW305-3-2-15-E-R2A2 as ProB, LRK54_RS01680 of *R. denitrificans* FW104-10B01 as TrpA, and BSU_32240 of *B. subtilis* 168 as ThrB). Finally, broadly speaking our assay identifies proteins *capable* of completing the missing function, which give significant insight into their own function. However, because of the uniqueness of overexpression in a heterologous host, compared to a potentially lower or context dependent expression in a native host, it is possible that this assay uncovers non-native function, and the putative cystathionine gamma-lyase from *R. denitrificans* FW104-10B01 found in the context of Δ*metB* may be an example.

## Discussion

Coaux-Seq provides another important tool for functional genomics and for advancing the annotation of genetic function. By creating DNA-barcoded overexpression libraries from randomly sheared genomic fragments, mapped using our established long-read sequencing workflow (Huang *et al*, 2022), we were able to repeatedly assay for the ability of contained genes to complete missing biochemical function. This was conducted in a cost-effective manner by utilizing BarSeq, which leverages primers containing Illumina adapters and only requires short sequencing read length. The Coaux-Seq assay not only provided the first experimental validation for the activity of 53 proteins, but also uncovered novel function.

Having established this workflow, it could readily be extended and in multiple ways. First, additional genomic libraries could be generated for newly isolated species. Second, additional genetic knock out backgrounds (auxotrophies) could be tested in *E. coli*, or in different species that may have different expression preferences (e.g., an alphaproteobacteria), although this may require a different expression vector. Alternatively, expression hosts like *E. coli* could be modified to improve the expression of heterologous DNA such as through provision of rare tRNAs (Cheng *et al*, 2015). Additional auxotrophies utilized could include genetic contexts that extend beyond single gene knockouts to target desired biochemical activity. Schema besides auxotrophies could also be used. Third, this assay could be extended to other sources of DNA beyond that of isolated bacterial genomes, such as field derived metagenomic DNA. However, this would require a more efficient assembly method for the barcoded libraries beyond blunt end ligation, such as one based on tagmentation (Crofts *et al*, 2021), to improve library diversity. All of these future assays would be benefited by improved transformation efficiency and improved means of high-throughput, particularly liquid, growth assays that do not bias the libraries to a few top candidates. Importantly, we have highlighted several ways in which one might observe false positives and false negatives in our complementation assay workflow for gene identification, providing a template for future users of such a method. Ultimately, Coaux-Seq provides a useful advance in functional genomics and has the potential to play an important role in future genetic annotation, particularly for core metabolism among non-model microbes. Improvements in such annotation stand to benefit downstream modeling approaches, including genome-scale metabolic models, with application in monoculture and microbial community contexts.

## Acknowledgements

We acknowledge the support of the Dean A. Richard Newton Memorial Professor Chair Funds. This work was supported by ENIGMA – Ecosystems and Networks Integrated with Genes and Molecular Assemblies (http://enigma.lbl.gov), a Science Focus Area Program at Lawrence Berkeley National Laboratory, supported by the United States Department of Energy, Office of Science, Office of Biological & Environmental Research under contract number DE-AC02-05CH11231. We acknowledge QB3 Genomics, University of California Berkeley (Berkeley, CA) for sequencing support through RRID:SCR_022170. This work was supported by NIH S10 OD018174 Instrumentation Grant.

## Author contributions

B.W.B., M.N.P., A.M.D., and A.P.A. conceived the project and wrote the manuscript. B.W.B., D.L., J.E., L.F., Y.Y.H. K.K., V.V.T., J.V.K., L.M.L., R.C., A.M.D. performed experiments and collected data. B.W.B., M.N.P., Y.Y.H., A.M.D., and A.P.A. analyzed data.

## Materials and Methods

### Culturing and media

Cultivations were incubated in a New Brunswick Innova 42 incubator. General cultivations were carried out in LB Miller medium at 37°C. For liquid cultures, 200 rpm shaking was used. After transformation, cells were recovered in either New England Biolabs (NEB) SOC or 10-β/stable recovery medium. Complementation selections were carried out in Difco M9 medium with calcium and magnesium added, and with 1% glucose as a carbon source. All agar plates were 1.5% agar (15 g agar per 1 L). Kanamycin, used for *Escherichia coli* Keio single gene knockout strain selection, was used at a concentration of 25 μg/mL. Kanamycin was prepared as 1000x stocks in water, sterile filtered (0.2 μm), and stored as 1mL aliquots in sterile 1.5 mL Eppendorf tubes at - 20°C. Chloramphenicol, which was used for plasmid selection, was used at a concentration of 17 μg/mL. Chloramphenicol was prepared as 500x stocks in 95% ethanol, sterile filtered (0.2 μm), and stored as 1 mL aliquots in sterile 1.5 mL Eppendorf tubes at −20°C. Anhydrotetracycline (aTc), used for synthetic promoter (tet) induction, was primarily used at 100 μg/mL (“1x”) unless otherwise noted. aTc was first prepared as a 100,000x stock in DMSO, the diluted to 1000x stocks in DMSO as needed. aTc stocks, with in a sterile 50 mL Falcon tube or a sterile 1.5 mL Eppendorf tube, were wrapped in aluminum foil to protect from light degradation and stored at −20°C.

### Cloning, strains isolation and genomic sequencing

Genome fragment library generation cloning was carried out in New England Biolabs 10-β high-efficiency electrocompetent *E. coli* cells. Individual knock out variants of *Escherichia coli* BW25113 for complementation transformations were generous gifts of Dr. Gareth Butland and were originally obtained from the Keio collection (Baba *et al*, 2006). *Sphingomonas koreensis* JSS26 DSMZ 15582 was obtained from DSM. *Escherichia coli* BW25113 wild-type was obtained from the Coli Genetic Stock Center at Yale University. *Bacillus subtills* 168 was obtained from the *Bacillus* Genetic Stock Center at The Ohio State University. *Bacteroides thetaiotaomicron* VPI-5482 was obtained from ATCC. *Pseudomonas fluorescens* FW300-N2E2, *Lysobacter sp.* FW306-1B-D06B, *Xylophilus sp.* GW821-FHT01B05, *Rhodanobacter denitrificans* FW104-10B01, *Rhodoferax sp.* GW822-FHT02A01, *Pedobacter sp.* FW305-3-2-15-E-R2A2, and *Acidovorax sp.* FHTAMBA were isolated from the Oak Ridge Reservation Field Site.

The full description of the isolation for *P. fluorescens* FW300-N2E2 has been given previously (Price *et al*, 2018). The isolation for *R. denitrificans* FW104-10B01 has been described elsewhere as well (Carlson *et al*, 2019). *Lysobacter sp* FW306-1B-D06B was isolated at 25°C, aerobically, in R2A medium in 96 well carbon from sample FWB306-02. *Pedobacter sp.* FW305-3-2-15-E-R2A2 was isolated aerobically in R2A medium at 30°C from sample FW305-03-02-15. *Xylophilus sp.* GW821-FHT01B05 was isolated aerobically in R2A medium at 25°C from sample GW821-2019-05-13-UF-R10. *Rhodoferax sp.* GW822-FHT02A01 was isolated using ground water pre-incubated at 4°C aerobically in R2A medium at 25°C from the GW822E-2019-04-01-UF-R10 sample as part of a high-throughput campaign. *Acidovorax sp.* FHTAMBA was isolated as part of high-throughput isolations from the Oak Ridge site. Previously sequenced genomes were available prior to this study for *E. coli* BW25113 (Grenier *et al*, 2014), *B. thetaiotaomicron* VPI-5482, *B. subtilis* 168, *P. fluorescens* FW300-N2E2 (Carim *et al*, 2021), *S. koreensis* JSS26, and *R. denitrificans* FW104-10B01 (Peng *et al*, 2022). For all other strains, the genomes were sequenced. For *Acidovorax sp*. FHTAMBA, *Xylophilus sp.* GW821-FHT01B05, *Lysobacter sp*. FW306-1B-D06B, and *Pedobacter sp.* FW305-3-2-15-E-R2A2, Plasmidsaurus (Oregon, USA) hybrid Nanopore/Illumina sequencing and assembly services were utilized. For *Rhodoferax sp*. GW822-FHT02A01 HMW DNA was extracted using the Genomic Tip 100/G kit. Nanopore and Illumina libraries were prepped in-house and sequenced as described in Goff et al 2022 (Goff *et al*, 2022). Sequence quality control is also the same as in Goff et al. 2022. The genome was hybrid assembled using Unicycler v0.4.8 (Wick *et al*, 2017) with default parameters. Each newly sequenced genome has been deposited to NCBI. The full set of accession number for genomes used in this study include, CP009273.1/GCF_000750555.1 (*Escherichia coli* BW25113), NZ_CP015225.1/GCF_001623525.1 (*Pseudomonas fluorescens* FW300-N2E2), GCF_002797435.1 (*Sphingomonas koreensis* JSS26; DSMZ 15582), NC_000964.3/GCF_000009045.1 (*Bacillus subtilis* 168), NC_004663.1/GCF_000011065.1 (*Bacteroides thetaiotaomicron* VPI-5482), CP088922.1/ NZ_CP088922.1/GCF_021560695.1 (*Rhodanobacter denitrificans* FW104-10B01), CP151802 (*Lysobacter sp*. FW306-1B-D06B), CP151803 (*Pedobacter sp*. FW305-3-2-15-E-R2A2), CP152408 (*Xylophilus sp*. GW821-FHT01B05), CP152407 (*Acidovorax sp*. FHTAMBA) and CP152376 (*Rhodoferax sp*. GW822-FHT02A01).

### General Cloning

The plasmid backbone pBbA2c-RFP (Addgene #35326) (Lee *et al*, 2011), a generous gift from Prof. Jay Keasling, was modified to generate the plasmids in this study. The initial three genome fragment libraries (*E. coli* BW25113, *Pseudomonas fluorescens* FW300-N2E2, *and Sphingomonas koreensis*) were created in the context of the pBbA2c-RFP derivative pBWB507, a variant of which containing a ribosomal binding site and mRFP in place of a gene fragment has been deposited to Addgene (Addgene #209325). A slight modification to this plasmid was made to include SapI type II restriction sites, pBWB514 (Addgene #209326). The additional restriction sites make possible the transfer of mapped libraries, meaning those with long read sequencing to connect the gene fragments to unique barcodes, to new vector backgrounds without the need for PCR, which could induce library bias. All of the other eight libraries were cloned into pBWB514. Both vectors contain mosaic ends (Crofts *et al*, 2021) to potentially allow for Gibson assembly cloning in future genome fragment library creation. Plasmid modifications were generated by successive rounds of PCR using PrimeSTAR by Takara following manufacturer protocols, (3 minutes 98°C denaturing, followed by 30x cycles of at a 10 second 98°C denaturing step, a 15 second 55°C annealing step, and a 45 second 72°C extension step, followed lastly by a 5 minutes 72°C extension). Self-ligation was used to create the new plasmids using NEB’s KLD kit, following manufacturer protocols. All primers used for plasmid modification, sequencing, and analysis are in the primer table among the supplemental materials.

### Genomic Fragment Library Preparation

Libraries were prepared by first growing the individual strains to an OD_600_ > 1.0 (37°C for *E. coli*; 30°C for all other strains). Cultures were carried out at 5 mL volume in 14 mL Falcon culture tubes. Genome extractions were carried out with a Thermo Scientific GeneJET Genomic DNA Purification kit, following manufacturer’s protocols, eluting into nuclease free water, and adjusting for gram-negative or gram-positive bacteria. Following extraction, genomes were sheared to a mean 3 kb fragment size with a Covaris S220. Between 2 and 20 μg of genomic DNA (typically 2 μg, total final volume 200 μL) was added into a Covaris blue miniTUBE, and the equipment was run using the 3kB setting (SonoLab 7.2 software, temperature set between 4 and 25°C, Peak Power 3.0, Duty Factor 20.0, Cycles/Burst 1000, 600 seconds). Following sonication, sheared genomes were run on a 1% agarose gel (100 volts, 400 mA, ∼25 minutes) and the “smeared” band between 1.5 and 5 kb was excised (Figure S12 for an example). Sheared genomic DNA was collected via gel extraction using a Thermo Scientific GeneJET Gel Purification kit, following manufacturer’s instructions but eluting into nuclease free water.

After gel extraction, genomic fragments were end-repaired using a Thermo Scientific Fast DNA End Repair Kit (manufacturer’s protocols). Between 2 and 4x 50 μL reactions were run and subsequently pooled using a Thermo Scientific GeneJET PCR Purification Kit (manufacturer’s protocols, nuclease free water elution). Even though the Fast DNA End Repair Kit phosphorylates the end-repaired DNA fragments, for thoroughness, an additional phosphorylation was conducted with NEB T4 Polynucleotide Kinase (PNK), following manufacturer’s protocols and using the 1X T4 DNA Ligase Buffer that contains 1 mM ATP. Multiple 50 μL reactions were run to include all end repaired DNA. After phosphorylation all reactions were again pooled using the Thermo Scientific GeneJET PCR purification Kit and eluted with 20 μL of nuclease free water.

Backbone was prepared by PCR to introduce barcode diversity via primer overhang. For pBWB507, to linearize and added the barcode and flanking regions used for BarSeq (described below) the forward primer used was BWB1429 (GTCGACCTGCAGCGTACGNNNNNNNNNNNNNNNNNNNNAGAGACCTCGTGGACATCactgtcTAGGGATAACAGGG) and the reverse primer was BWB1422 (atggtctgaattcttttctctatcac). Similarly, for pBWB514, BWB1471 (CTGTCTCTTATACACATCTGTCGACCTGCAGCGTACGNNNNNNNNNNNNNNNNNNNNAGAGACCTCGTGGACATCG) and BWB1472 (CTGTCTCTTATACACATCTTGGACTGAAGAGCttttctctatc) were used. All primers were ordered from IDT. None of these primers were phosphorylated and BWB1429 and BWB1471 were HPLC purified to help ensure full length products from PCR. Backbone preparation was run in 4x of 50 μL PCR using PrimeSTAR Max from Takara as described above. Following PCR, 1 μL of DpnI was added to each PCR reaction and the reaction was incubated at 37°C for 1 hour to remove the PCR template plasmid. Following, a 1% agarose gel was run and the ∼3 kb backbone was excised and purified by gel extraction. Out of an abundance of caution, an NEB rSAP reaction was run on the purified backbone to ensure no phosphorylation of the backbone to avoid self-ligation during blunt-end ligation cloning. After the rSAP reaction, another GeneJET PCR purification was run.

### Blunt end ligation, genome fragment library transformation

Once both insert and backbone DNA elements were prepared as described above, blunt end ligation with NEB T4 DNA ligase was conducted (2,000,000 units/mL). The ligations were carried out at a 1:3 backbone to insert molar ratio, run in 4x at a 20 μL volume, and reactions were run at 16°C overnight (∼16 hour). After overnight incubation, the 4x ligations were pooled via a GeneJET PCR Cleanup Kit and the DNA concentration was measured with a Thermo Scientific NanoDrop 2000 using the NanoDrop 2000 software. Following, 3-5 μL of this pooled ligation product was added to NEB 10-β high-efficiency electrocompetent cells (25 μL) in a VWR 90 μL sterile electroporation cuvette and were electroporated with a BTX Harvard Apparatus Electro Cell Manipulator PrecisionPulse (ECM 630, Version 1.05) with a BTX Harvard Apparatus Safety Stand (630B). Electroporation settings were 1750 volts, 200 Ω resistance, and 25 μF capacitance. Immediately after electroporation, cells were recovered with 975 μL of 10-β/stable NEB recovery medium for 1 hour at 37°C and subsequently stored at 4°C until later use. To ensure sufficient library size, up to 4 separate ligation product transformations were conducted per library.

### Plating, library outgrowth

To determine transformation efficiency, 10-fold serial dilutions in sterile 1x PBS were plated as 15 μL spots on LB agar chloramphenicol plates. After drying, plates were incubated at 37°C overnight. The next morning colonies were counted to determine the colony forming units per volume (cfu/mL), and thus determine the volume of recovered electroporation media necessary to target a generated a fragment library to sufficiently “cover” or sample the genome. Coverage need was estimated by taking the number of putative genes in the genome (typically 3,000 to 6,000) and the multiplying by ∼10x (30,000-60,000 cfus, typically). This calculated volume of the recovered electroporation was then added to 50 mL of LB medium with chloramphenicol and allowed to grow overnight in a 250 mL Erlenmeyer flask. The next morning, 4x glycerol stocks (1 mL volume equal parts 50% glycerol and culture medium) were prepared and 6x 5 mL aliquots of the culture medium were used for a miniprep to recovery the plasmid library using a Thermo Scientific GeneJET Plasmid Miniprep Kit, following manufacturer’s instruction but eluting with nuclease free water. Cultures aliquoted for the minipreps were centrifuged for 10 minutes at 4000 x *g* at 4°C to pellet the cells. After decanting the media, remaining media was gently removed by pipetting. The plasmid library obtained from miniprep was subsequently used for BarSeq to determine library size by sequencing, PacBio sequencing to further validate library size and quality and to link gene fragment to barcodes, and complementation transformations.

### Preparation of Keio competent cells and electroporation

To prepare individual single gene knockout auxotrophic *E. coli* strains from the Keio collection for electroporation, they were first inoculated into 3 mL of LB with kanamycin and incubated overnight at 37°C, 200 rpm shaking. The following morning, the overnight culture was diluted 1:100 into 50 mL of fresh medium and grown at 37°C, 200 rpm shaking. The OD_600_ was measured until the cells reached an OD_600_ between 0.4-0.6. Following, the volume was split in half (∼25 mL), and centrifuged for 10 minutes at 4000 x *g* at 4°C to pellet the cells. All following steps were conducted on ice. After centrifugation, media was decanted and the cell pellet was resuspended in 5 mL of 10% glycerol that had been stored at 4°C and kept on ice. Cells were pipetted gently to resuspend fully. After resuspension, cells were spun down again at 4000 x g at 4°C for 10 minutes. This 10% glycerol washing process was repeated 3 additional times. After the 4^th^ wash, the cell pellet was resuspended into 100^th^ of the original volume in 10% glycerol and stored at 30 μL aliquots in sterile 1.5 mL Eppendorf tubes and either stored at −80°C or used immediately for transformation.

Electroporation was accomplished by adding 75 ng of plasmid DNA (3-5 μL volume) to the 30 μL competent cell aliquot, mixing by gently flicking the tube, transferring to an electroporation cuvette, and electroporating. After electroporating with a BTX Electro Cell Manipulator (PrecisionPulse) at 1750 V, 200 ohms resistance, and 25 μF capacitance, 975 μL of recovery medium was immediately added. For library recovery, LB medium was used. For selective recovery, complete M9 medium with 1% glucose was used. In either case, electroporated cells in the recovery medium were transferred to a 37°C shaker incubator for 1 hour. After 1 hour, the recovered cells were either made into a glycerol stock (mixed 1:1 with 30% glycerol to make a 15% glycerol stock solution) and frozen at −80°C, washed 3x in PBS before transfer to a selective condition (if in LB), or directly plated into a selective condition (if recovered in M9 medium).

### PacBio Sequencing

Preparation for long read sequencing began with 4x 50 μL PCR, using PrimeSTAR Max, as described above with the exception that 100 ng of plasmid library was used as template and with only 10x PCR cycles. For this PCR, both primers were HPLC purified and 5’ phosphorylated. For libraries on pBWB507 the primers were (Fw - /5Phos/gtgatagagaaaagaattcagaccat and Rv - /5Phos/cagtGATGTCCACGAGGTCTC) and for pBWB514 (Fw - /5Phos/GTCCAAGATGTGTATAAGAGACAG and Rv - /5Phos/GACGATGTCCACGAGGTCTC). After PCR, the product was run on 1% agarose gel and excised to gather the 1-5 kb range and pooled. After gel extraction, the PCR product was digested with DpnI at 37°C for 1 hour. After digestion, reactions were pooled using a GeneJET PCR purification. From this point, the PCR products of the DNA fragment libraries were prepared for PacBio sequencing using the PacBio SMRTbell prep kit 3.0, following the adapter-barcoded protocol (Barcoded overhang adapter kits 8A and 8B) including the initial bead clean up. Barcoded libraries were pooled and submitted for sequencing with the University of California-Berkeley QB3 Genomics core and sequenced with their PacBio Sequel II (SMRT cell 8M).

### Computational pipeline used to map barcodes to genes

The Boba-seq script was used to map barcodes to genomic inserts and to genes and can be found at https://github.com/OGalOz/Boba-seq (Huang *et al*, 2022). Briefly, assembly sequence files (FASTA) and annotation files (GFF/GFF3) for each source isolate were either downloaded from RefSeq or generated as part of sequencing work in this study. Default parameters were used for all steps as listed on the configuration .JSON file. PacBio’s lima tool, usearch (www.drive5.com/usearch/), vsearch (https://github.com/torognes/vsearch), and minimap2 (Li, 2018) are used as part of the pipeline. First, PacBio CCS reads are demultiplexed and then the insert and barcode sequences are extracted. Reads with concatemers or with incorrect barcode lengths are filtered out. Only inserts with 10 or fewer expected errors are kept. Inserts are then mapped to the genome assembly and a series of criteria are used to identify high-confidence mappings. Finally, genomic positions are mapped to protein-coding genes to generate final mapping tables used to calculate library statistics and to identify gene hits from complementation assays.

### Rare codon analysis

We computed the codon adaptation index (CAI) for each gene using the CAI program from EMBOSS (Rice *et al*, 2000) 6.6.0 with their reference table of codon usage in highly-expressed genes of *E. coli*. We computed the frequency of codons that are rare in *E. coli* (ATA, CGG, CGA, CTA, AGA, AGG, GGA, or CCC) in each gene that was expected to complement one of the Keio knockouts. There was no significant difference in the total frequency of rare codons between expected hits and those that were not hits.

### BarSeq

Barcodes were amplified using “BarSeq_V4” primers, as described previously (Huang *et al*, 2022). These use a 10-bp index sequence for P7 primers and an internal 8-bp index (to detect index hopping) and allow multiplexing up to 768 samples. The P7 BarSeq primers support demultiplexing by Illumina software via index reads, while the additional index added by the P5 primers is checked by the MultiCodes.pl script. BarSeq reads were converted to counts per barcode using the MultiCodes.pl script in the feba code base (https://bitbucket.org/berkeleylab/feba/), with the `-minQuality 0` option and `-bs4` (BarSeq 4) for BarSeq primers. To estimate the diversity of barcoded libraries, only reads that had a quality score of ≥30 at each position (the `-minQuality 30` option) were used, which corresponds to an error rate for barcodes of at most 0.001 * 20 nt = 2%. Furthermore, any barcodes that were off-by-1 errors from a more common barcode were eliminated. Per-strain fitness and z-scores were computed using bobaseq.R (https://github.com/morgannprice/BobaseqFitness) (Huang *et al*, 2022). These R scripts were also used to identify hits that were confirmed by overlap.

