## Supplemental Materials for "High-throughput protein characterization by complementation using DNA barcoded fragment libraries"

Bradley W. Biggs[1], Morgan N. Price[1], Dexter Lai[2], Jasmine Escobedo[2], Luis Fortanel[2], Yolanda Y. Huang[1], Kyoungmin Kim[2], Valentine V. Trotter[1], Jennifer V. Kuehl[1], Lauren M. Lui[1], Romy Chakraborty[1], Adam M. Deutschbauer[1,3], Adam P. Arkin[1,2]*

1Environmental Genomics and Systems Biology Division, Lawrence Berkeley National Laboratory, Berkeley, CA 94720, USA
2Department of Bioengineering, University of California-Berkeley, Berkeley, CA 94720, USA
3Department of Plant and Microbial Biology, University of California-Berkeley, Berkeley, CA 94720, USA

### Table of Contents

Supplemental Figures…………………………………………………………………………………….. pg. 3-19

Supplemental Tables……………………………………………………………………………………….. pg. 20-29

### Supplemental Figures

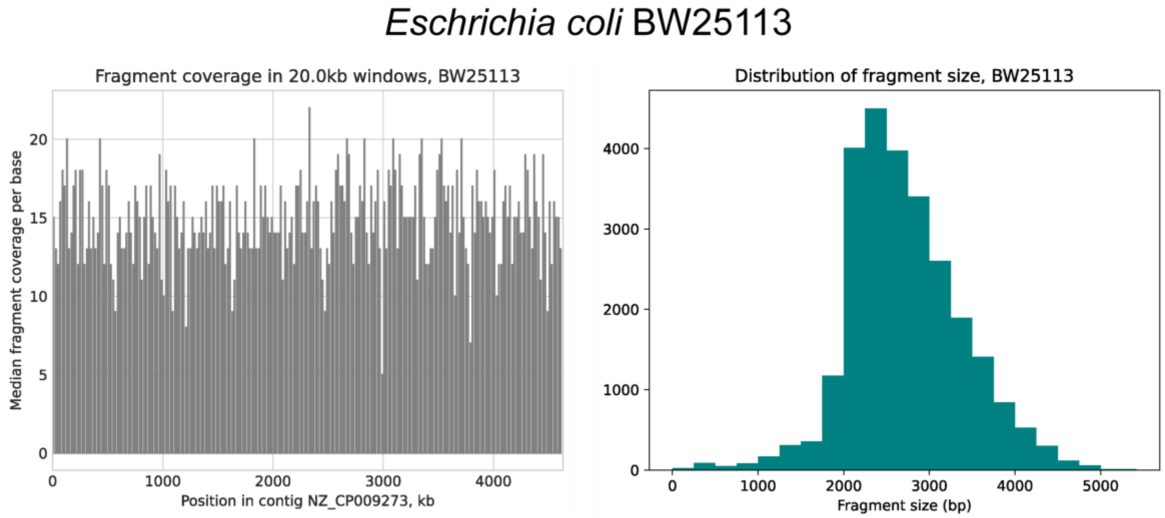


**Figure S1.** *Escherichia coli BW25113 fragment library quality figures. For Figure S1 through Figure S11, the left-hand side of the figure shows a plot of the fragment coverage of the genome. Specifically, the plot shows counts for the number of fragments that give coverage of a region of the genome, with a 20 kb window. The greater the height of the bar, the greater the number of fragments that cover that region of the genome. If there is no bar for a region, no fragment covered it. The plot on the right-hand side of the figure shows the distribution of fragment length. An average of 3 kb sheared genome fragments was sought, but all the distributions show an average below this, but with fragments both above and below the target size. The mean is typically closer to 2 kb than 3 kb.*

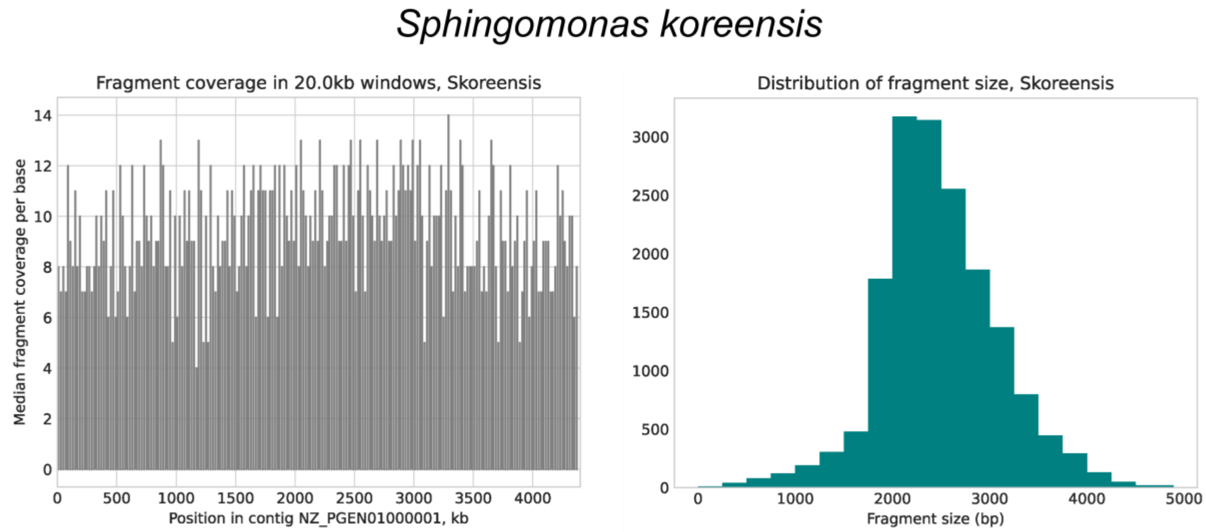


**Figure S2.** *Sphingomonas koreensis (JSS26, DSMZ 15582) fragment library quality figures.*

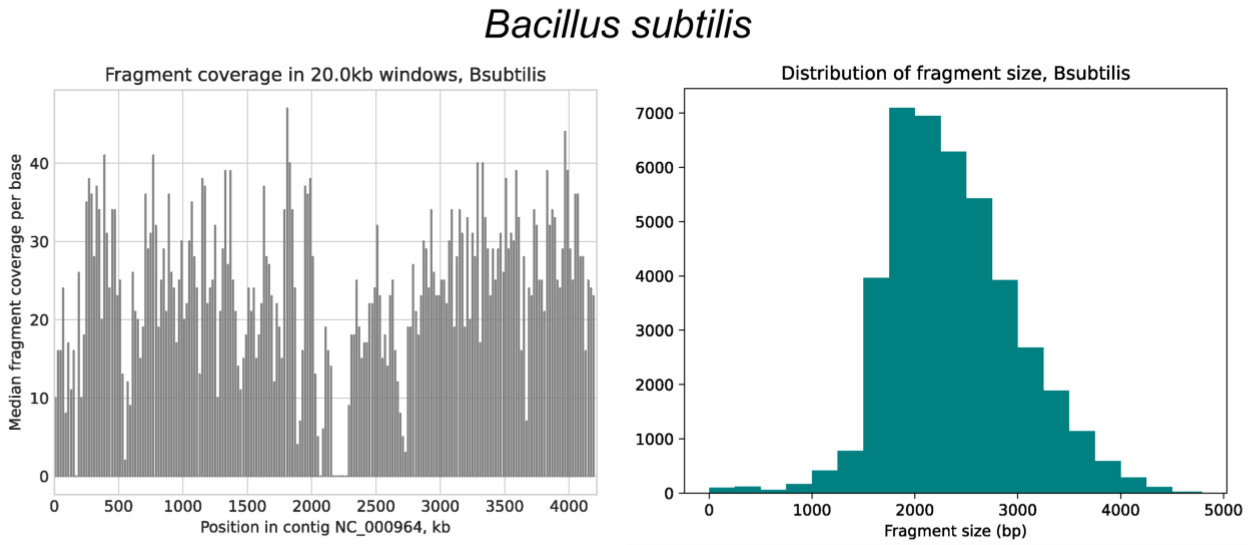


**Figure S3.** *Bacillus subtilis fragment library quality figures.*

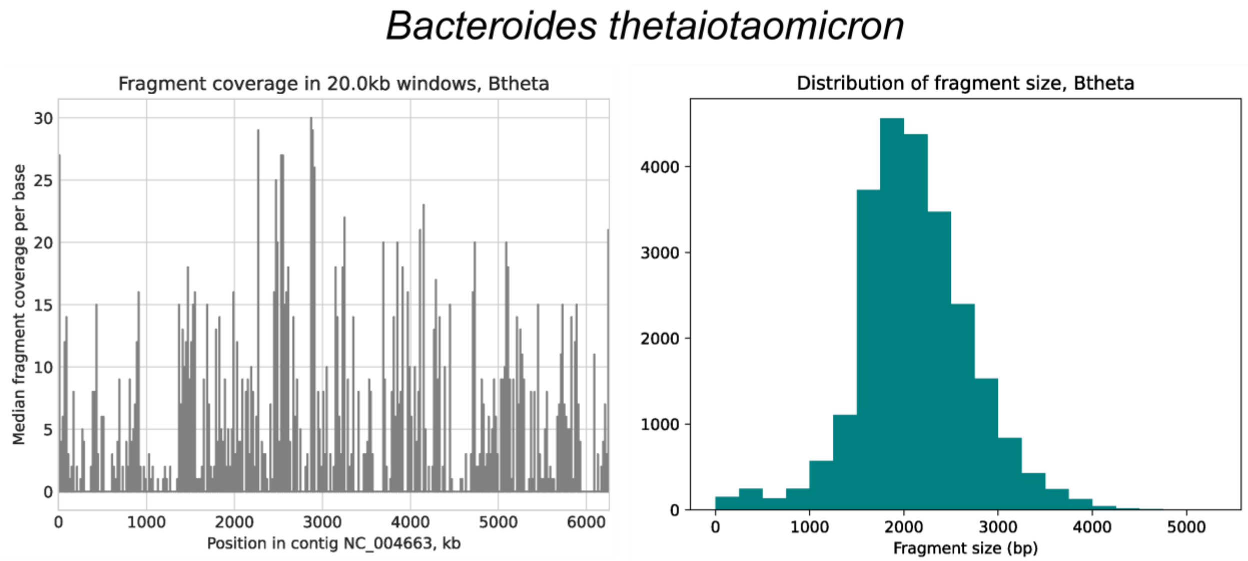


**Figure S4.** *Bacteroides thetaiotaomicron fragment library quality figures.*

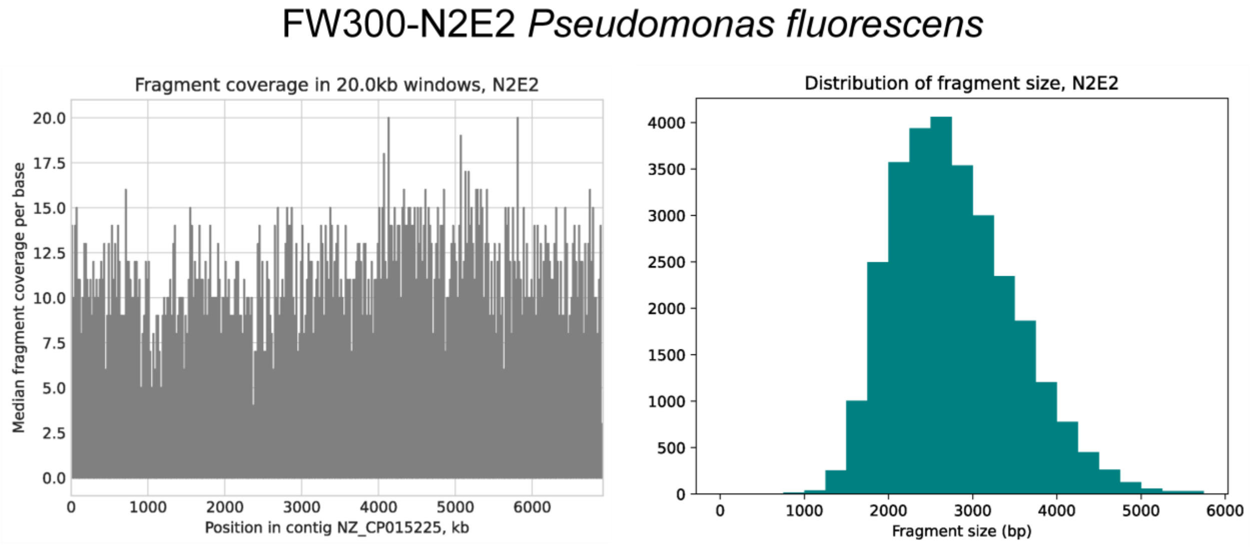


**Figure S5.** *FW300-N2E2 Pseudomonas fluorescens fragment library quality figures.*

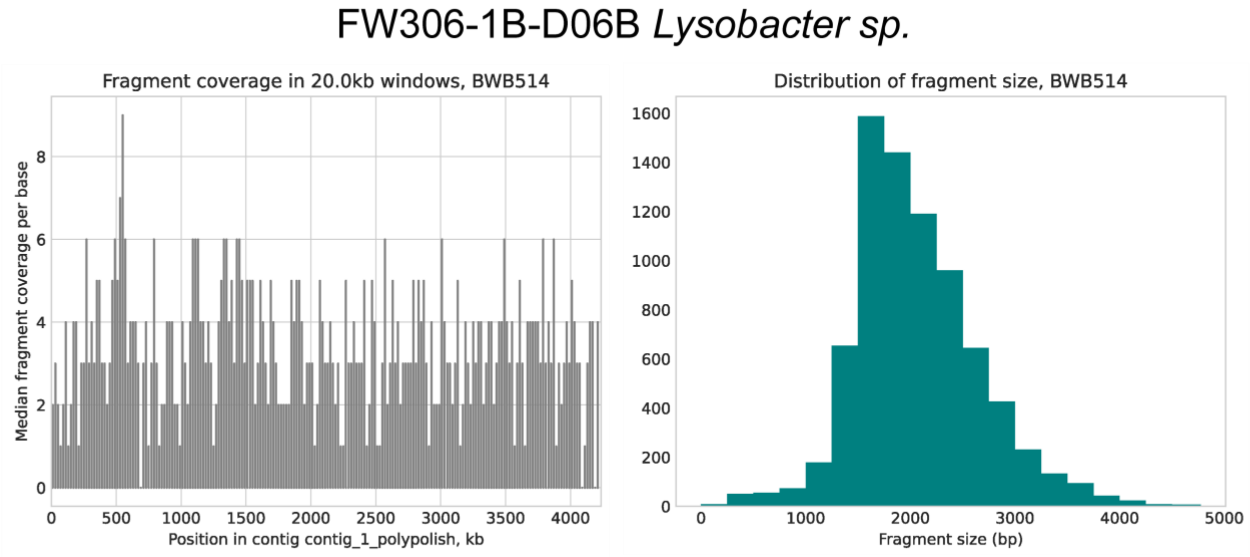


**Figure S6.** *FW306-1B-D06B Lysobacter sp. fragment library quality figures.*

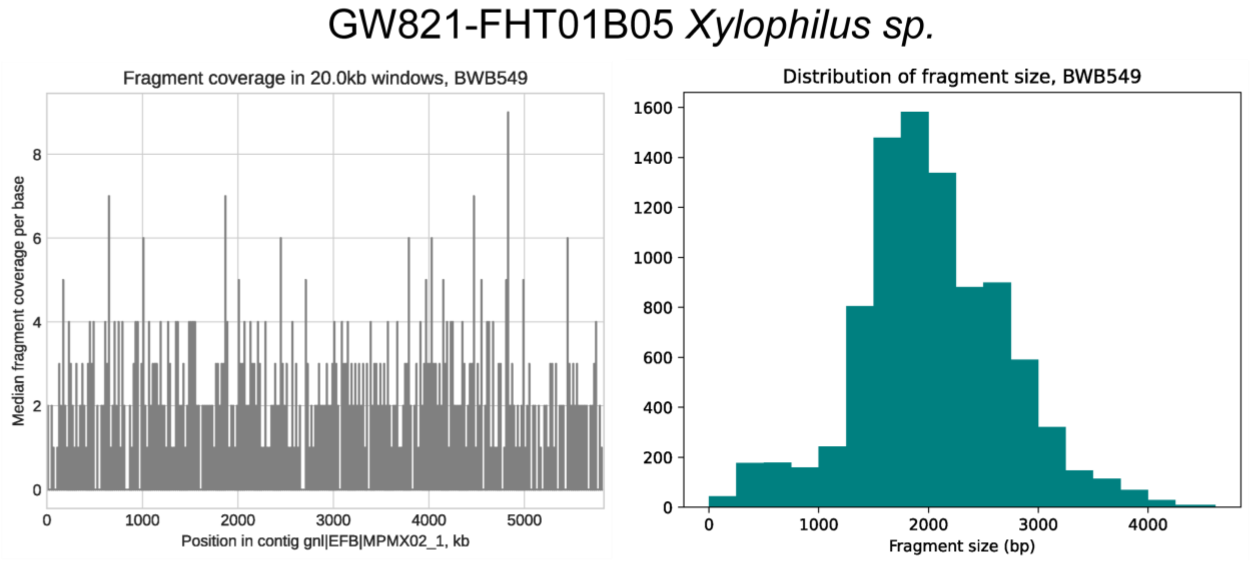


**Figure S7.** *GW821-FHT01B05 Xylophilus sp. fragment library quality figures.*

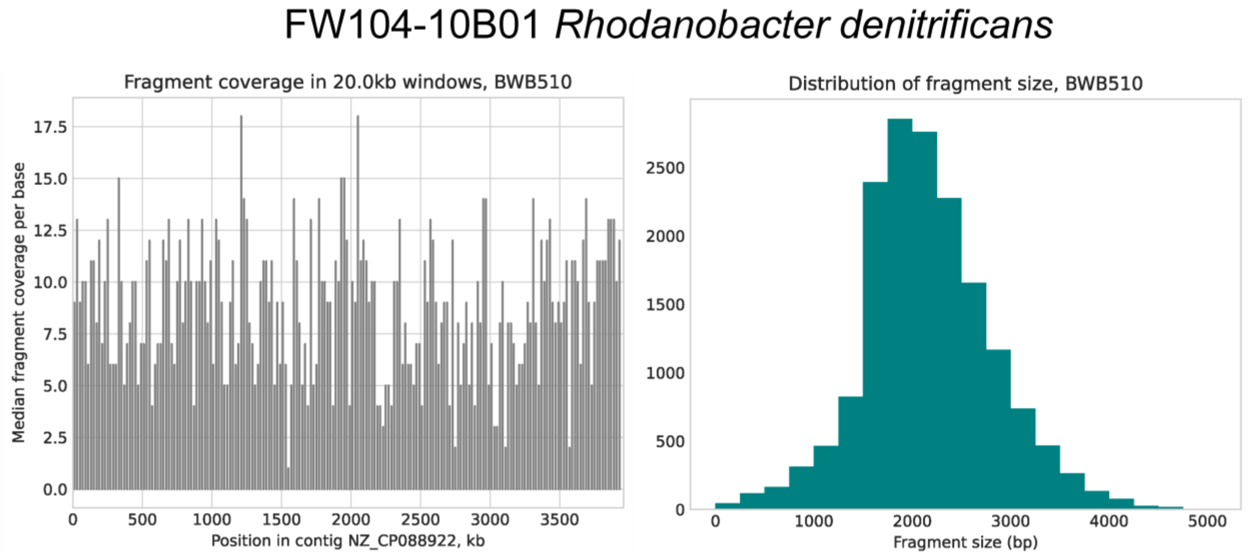


**Figure S8.** *FW104-10B01 Rhodanobacter denitrificans Fragment library quality figures.*

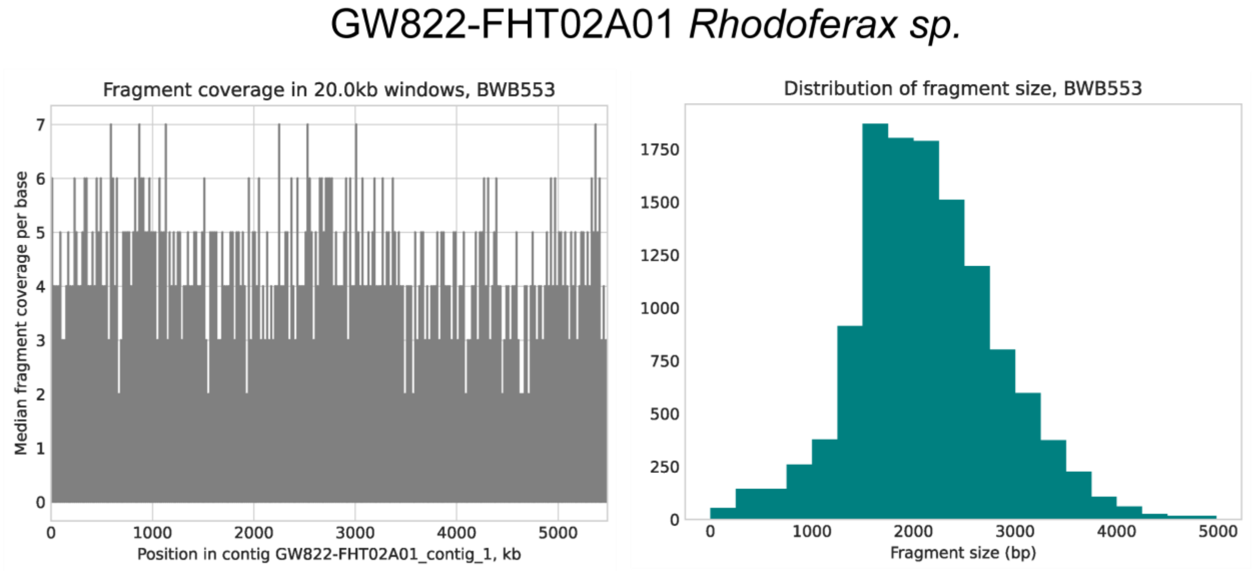


**Figure S9.** *GW822-FHT02A01 Rhodoferax sp. fragment library quality figures.*

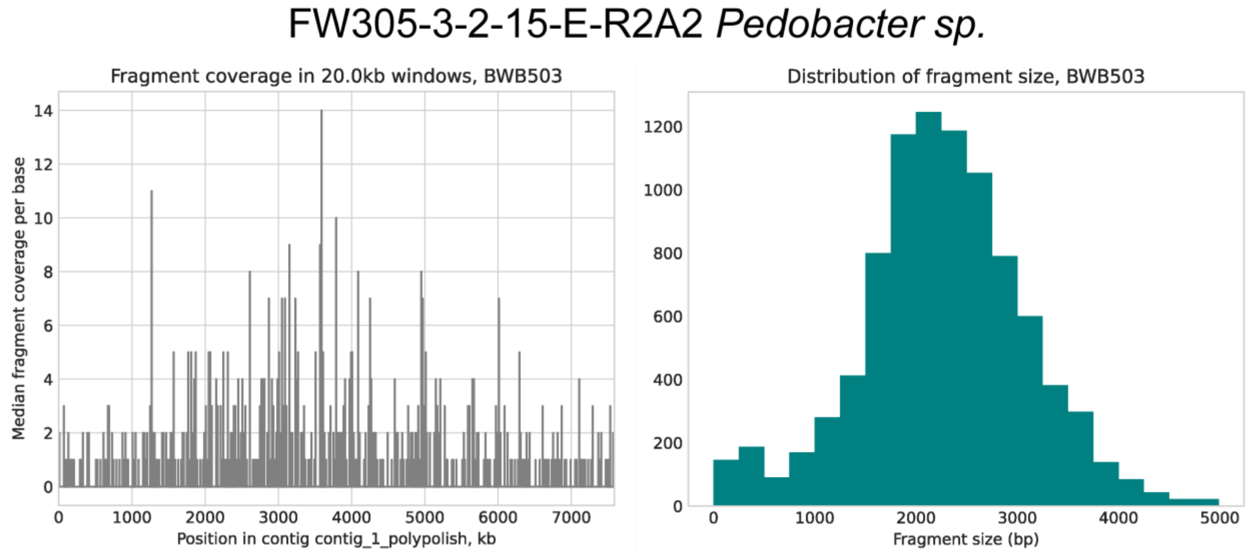


**Figure S10.** *FW305-3-2-15-E-R2A2 Pedobacter sp. fragment library quality figures.*

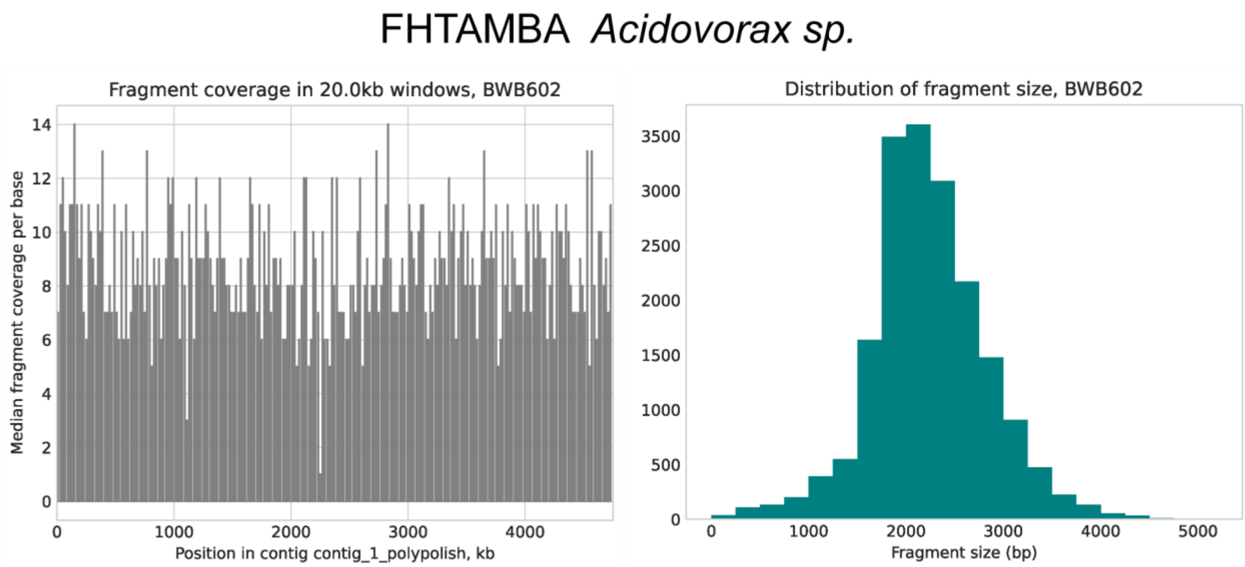


**Figure S11.** *FHTAMBA Acidovorax sp. Fragment library quality figures.*

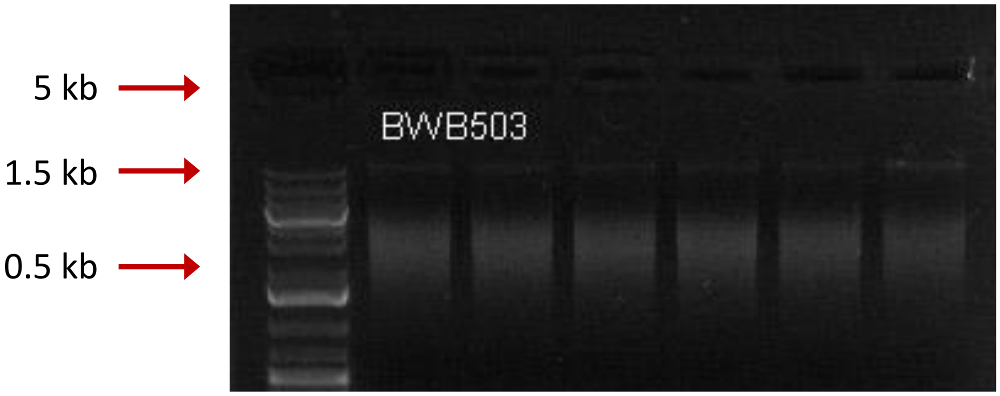


**Figure S12.** *Genome shearing example.* Example 1% DNA gel for Covaris ~3 kb shearing of 2 µg of genomic DNA from FHTAMBA (*Acidovorax sp.*). Ladder shown is Thermo GeneRuler 1 kb plus.

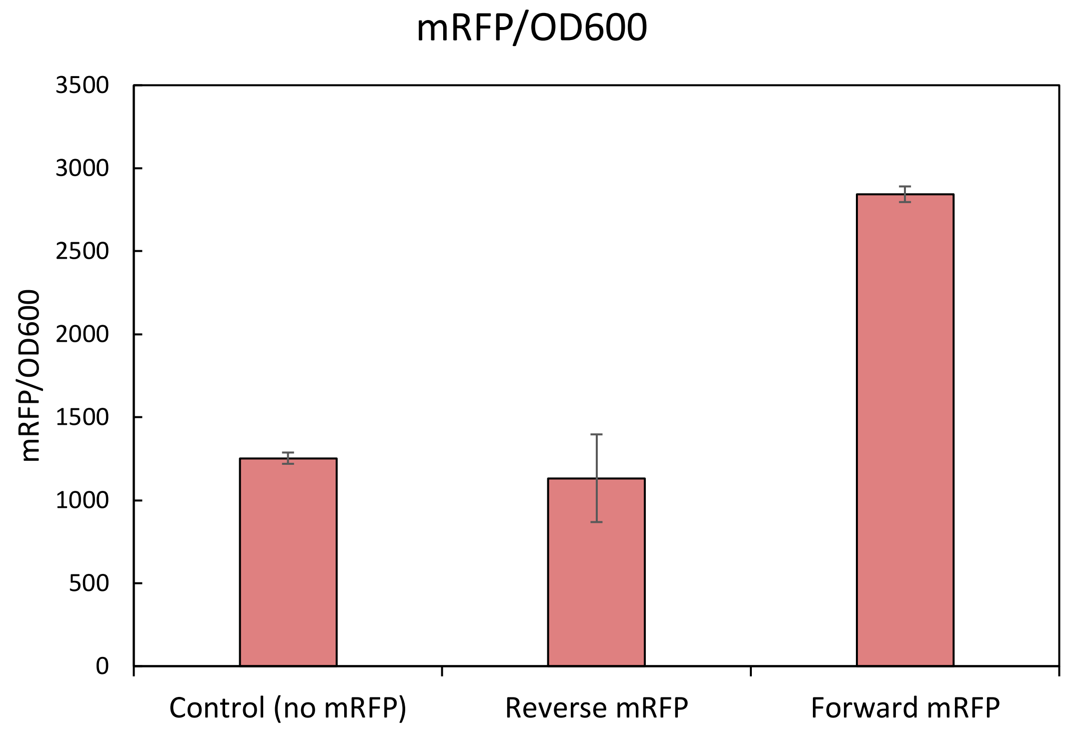


**Figure S13.** *mRFP expression confirmation.* Three constructs tested to confirm expression from the synthetic promoter in the context of the vector used in this study. All constructs contain a Tet promoter and were induced with 1x aTc. The control lacked any mRFP gene. "Reverse mRFP" had the mRFP gene in the reverse orientation with respect to the synthetic promoter. "Forward mRFP" possessed the mRFP gene in the correct orientation with the synthetic promoter. Fluorescence was read with excitation at 587 nm and emission at 610 nm. Emission values were divided by the $OD_{600}$ reading of the well. Values are for biological triplicate with the error bars representing standard deviation.

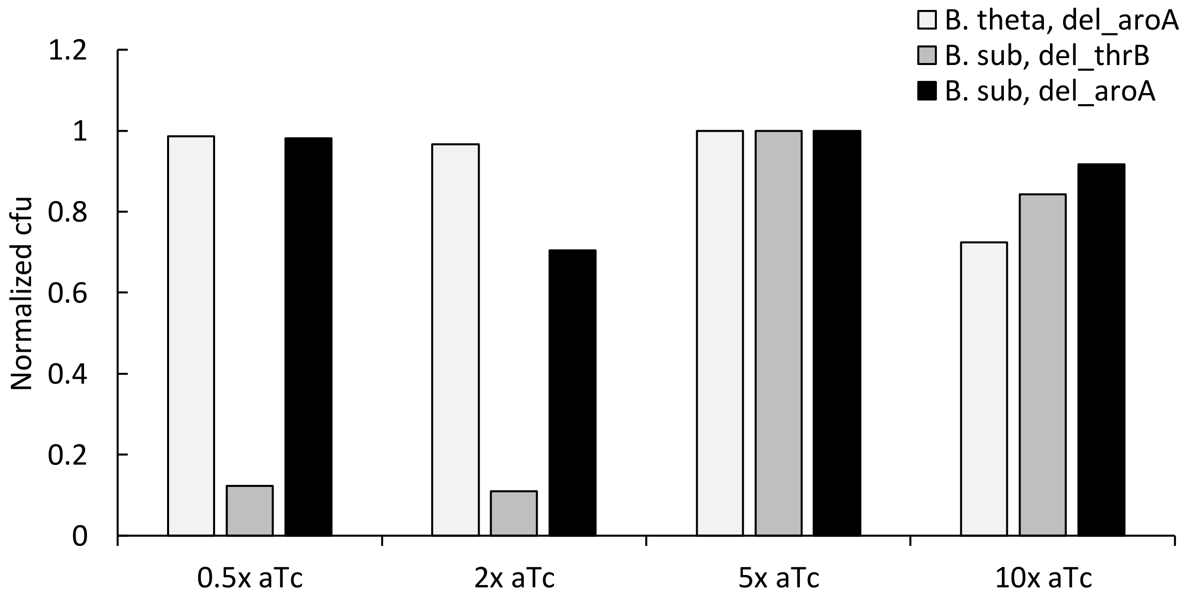


**Figure S14.** *Normalized colony forming units per aTc induction level for three complementation assays.*

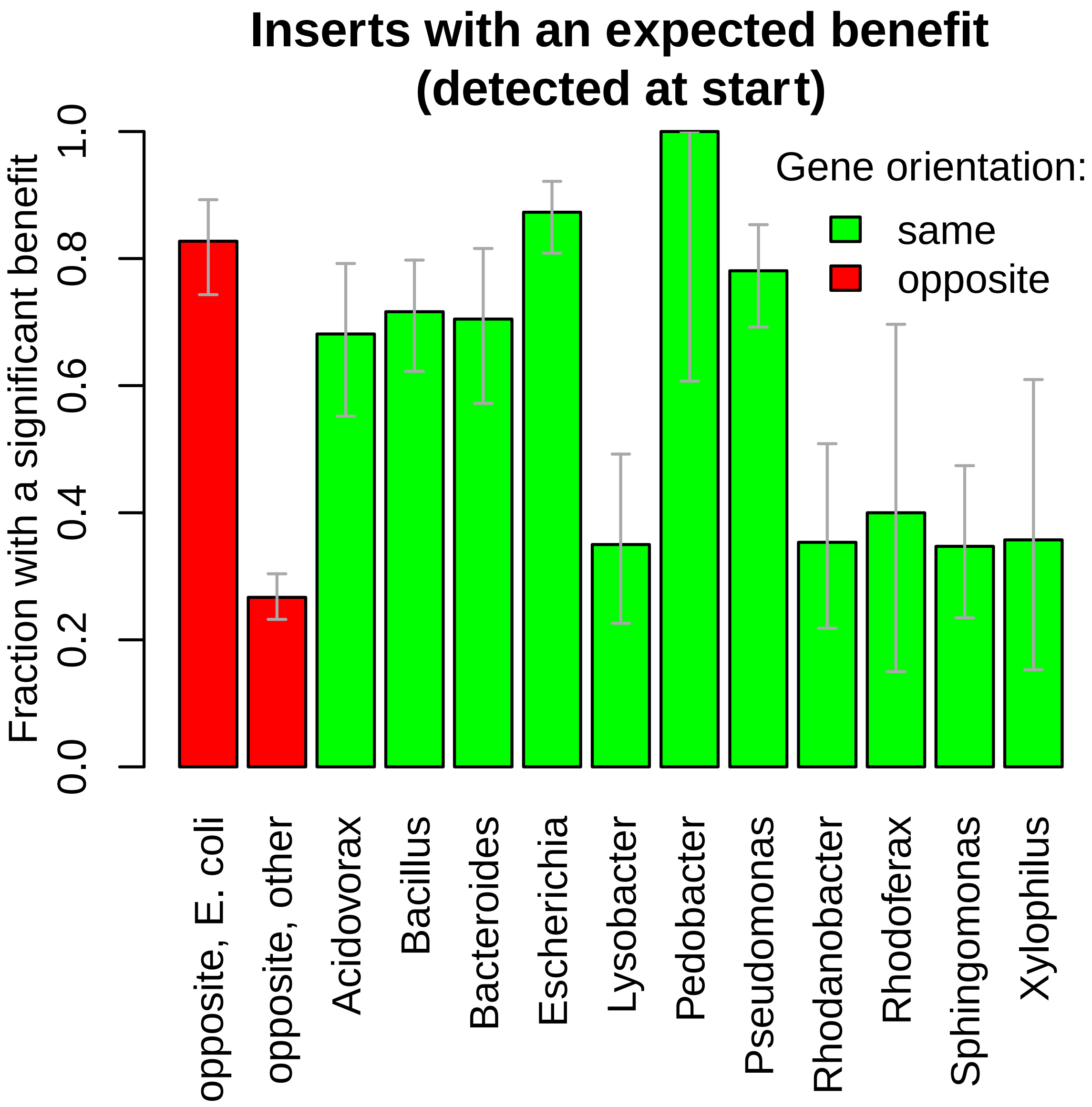


**Figure S15.** *Fraction of inserts that were expected to provide a benefit that do provide a high confidence benefit per the different genus of origin.*

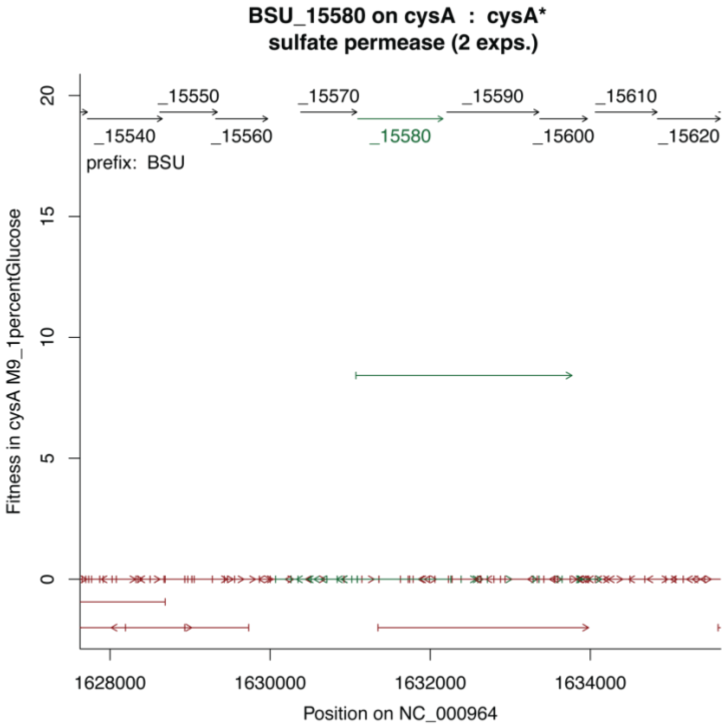


**Figure S16.** *Fitness benefit of BSU_15580 of B. subtilis associated fragments in the context of ΔcysA.*

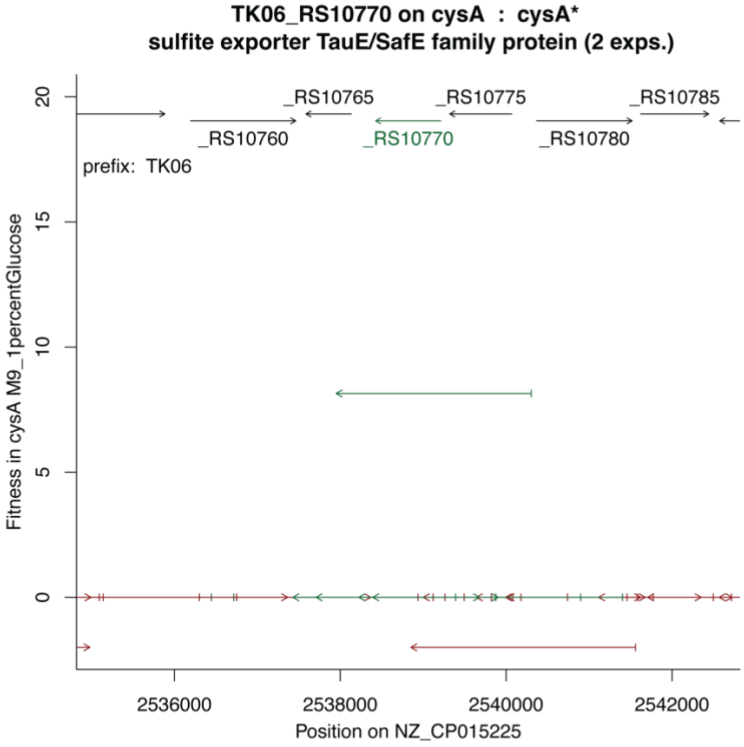


**Figure S17.** *Fitness benefit of TK06_RS10770 of P. fluorescens FW300-N2E2 associated fragments in the context of ΔcysA.*

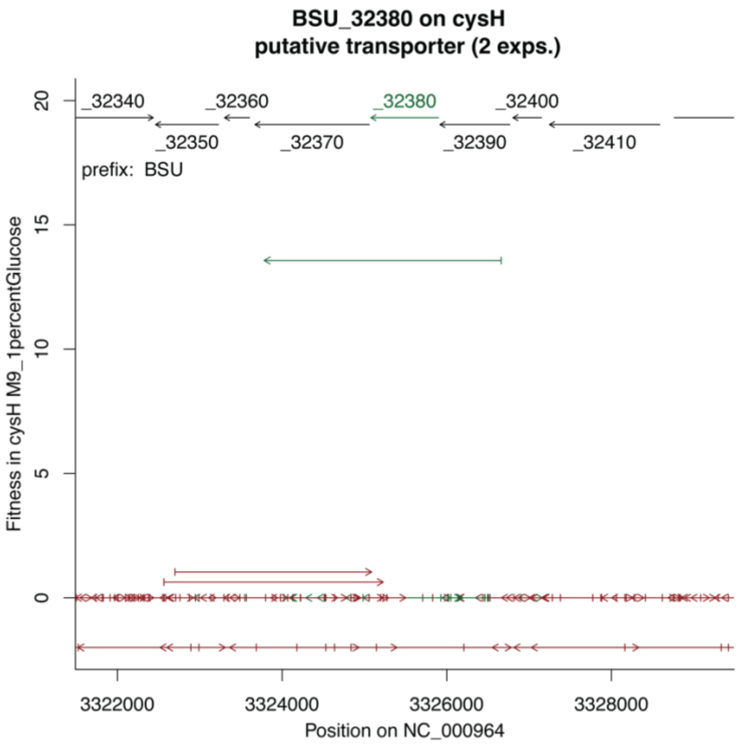


**Figure S18**. *Fitness benefit of B. subtilis BSU_32380 associated fragments in the context of ΔcysH.*

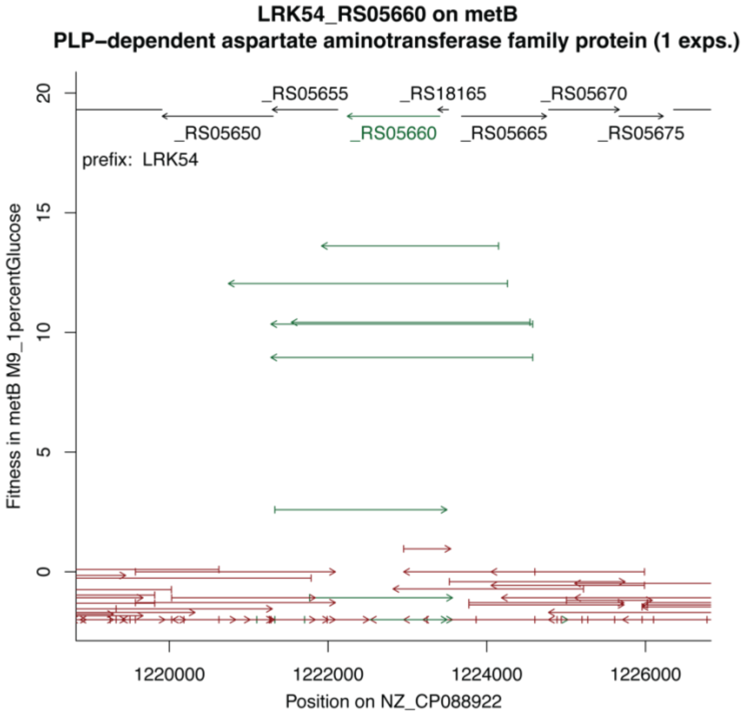


**Figure S19**. *Fitness benefit of LRK54_RS05660 of Rhodanobacter denitrificans FW104-10B01 associated fragments in the context of ΔmetB.*

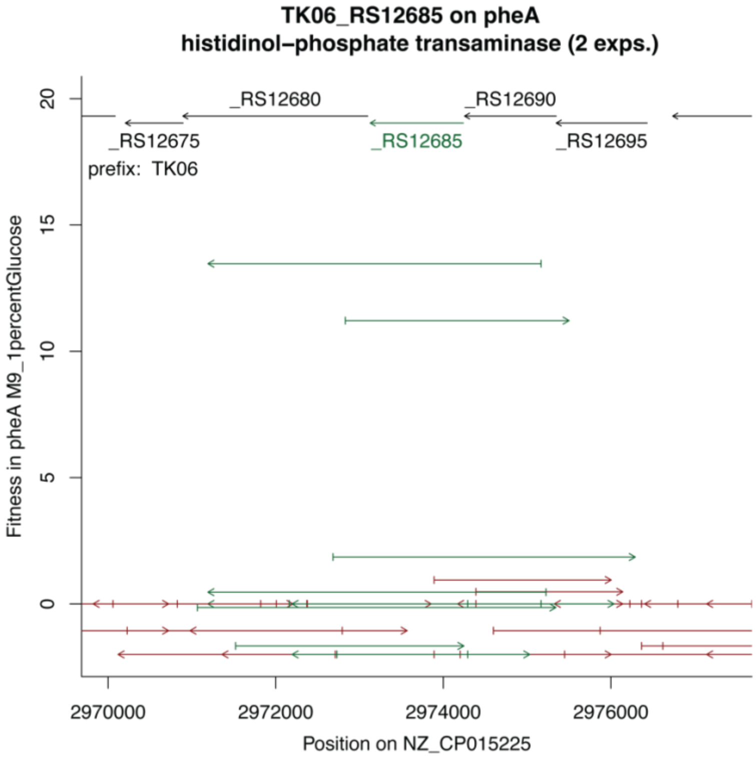


**Figure S20**. *Fitness benefit of TK06_RS12685 from P. fluorescens FW300-N2E2 associated fragments in the context of ΔpheA*.

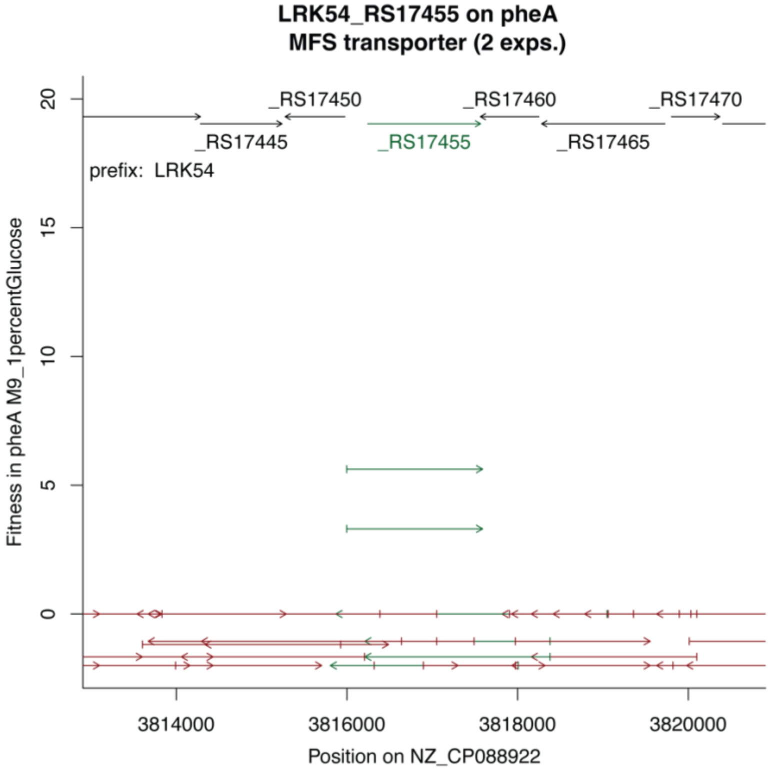


**Figure S21.** *Fitness benefit of LRK54_RS17455 of Rhodanobacter denitrificans FW104-10B01 associated fragments in the context of ΔpheA.*

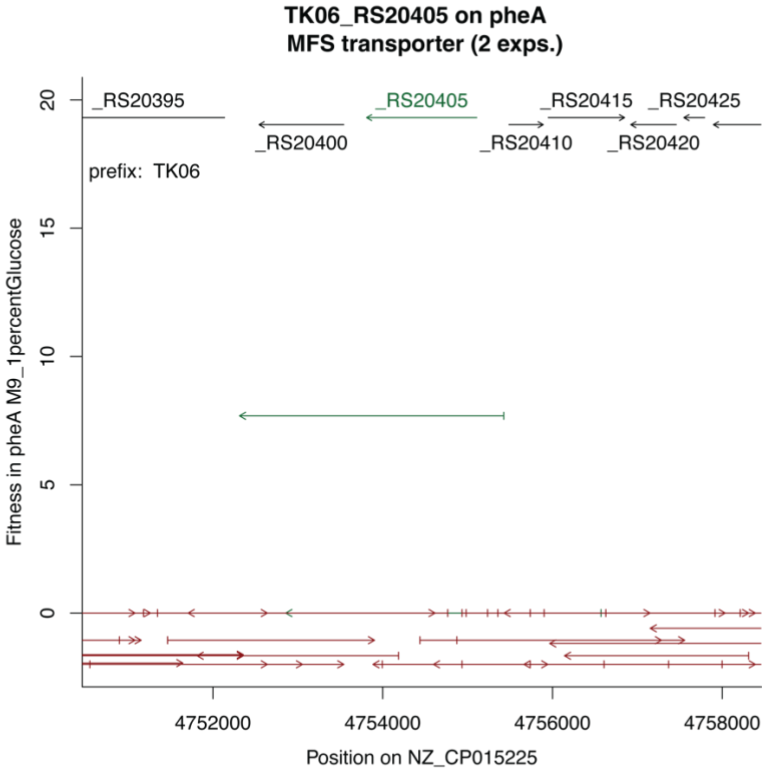


**Figure S22**. *Fitness benefit of TK06_RS20405 of P. fluorescens FW300-N2E2 associated fragments in the context of ΔpheA.*

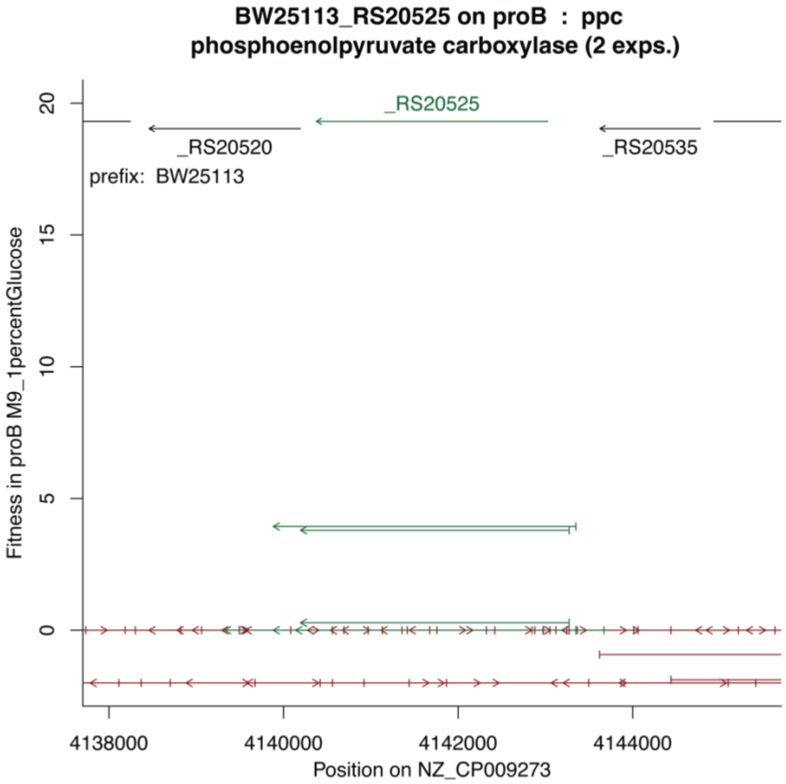


**Figure S23**. *Fitness benefit of E. coli BW25113_RS20525 associated fragments in the context of ΔproB.*

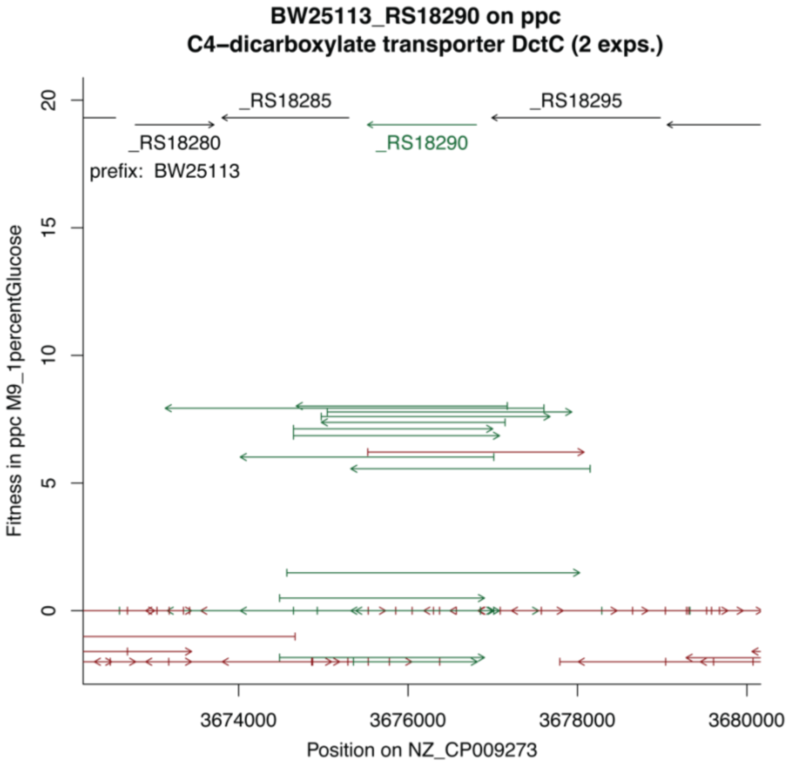


**Figure S24**. *Fitness benefit of E. coli BW25113_RS18290 associated fragments in the context of Δppc.*

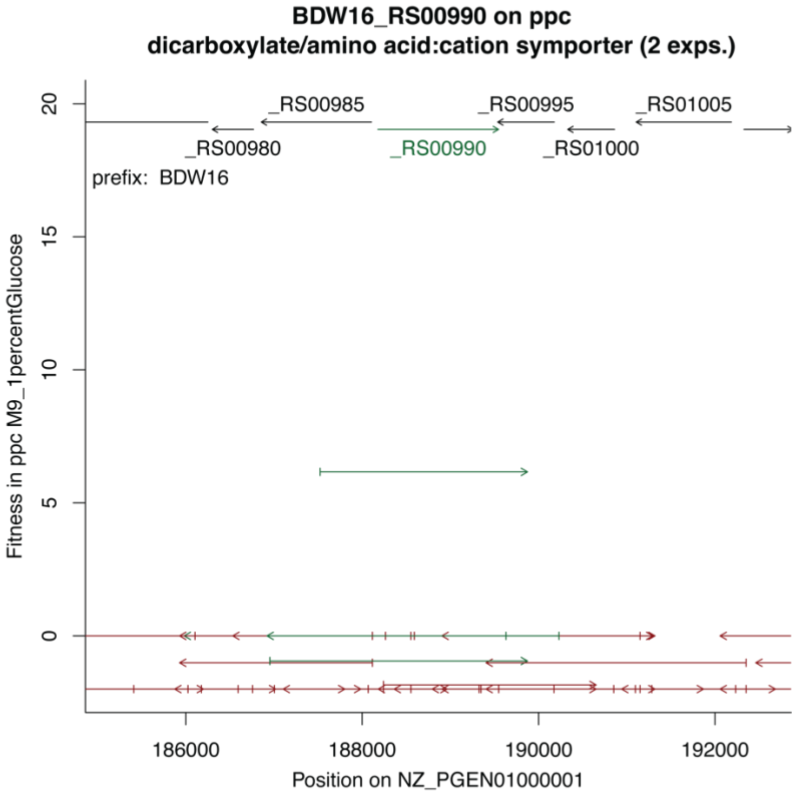


**Figure S25**. *Fitness benefit of BDW16_RS00990 of S. koreensis associated fragments in the context of Δppc.*

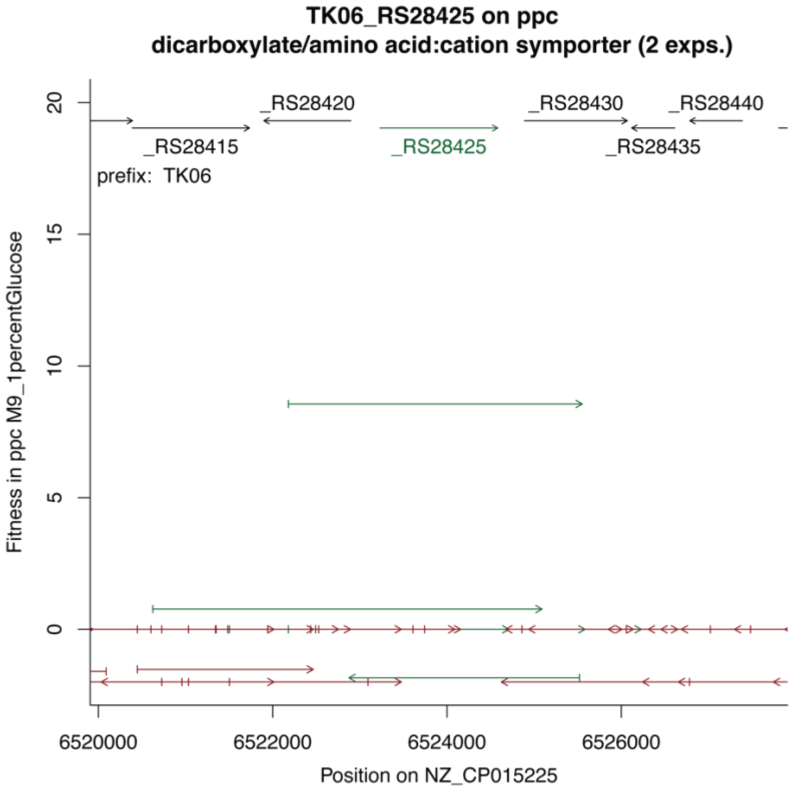


**Figure S26**. *Fitness benefit of TK06_RS28425 of P. fluorescens FW300-N2E2 associated fragments in the context of Δppc.*

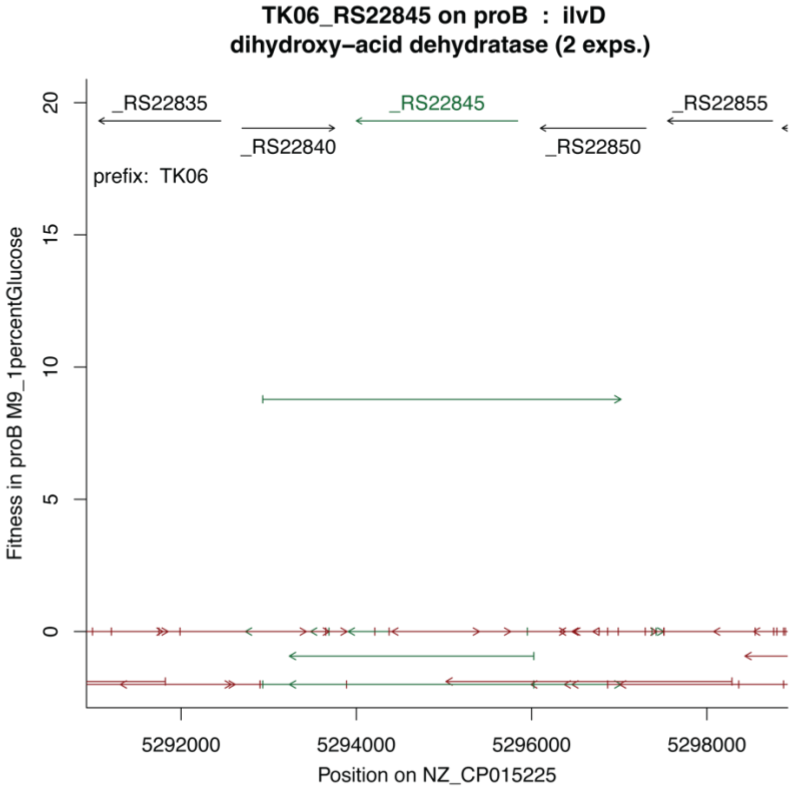


**Figure S27**. *Fitness benefit of TK06_RS22845 of P. fluorescens FW300-N2E2 associated fragments in the context of ΔproB.*

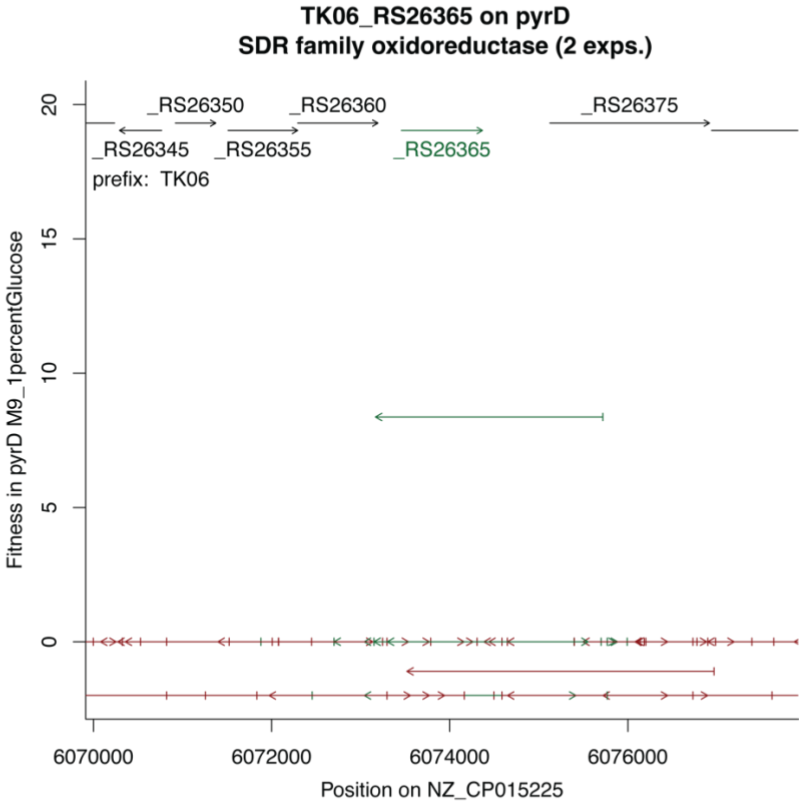


**Figure S28**. *Fitness benefit of TK06_RS26365 of P. fluorescens FW300-N2E2 associated fragments in the context of ΔpyrD.*

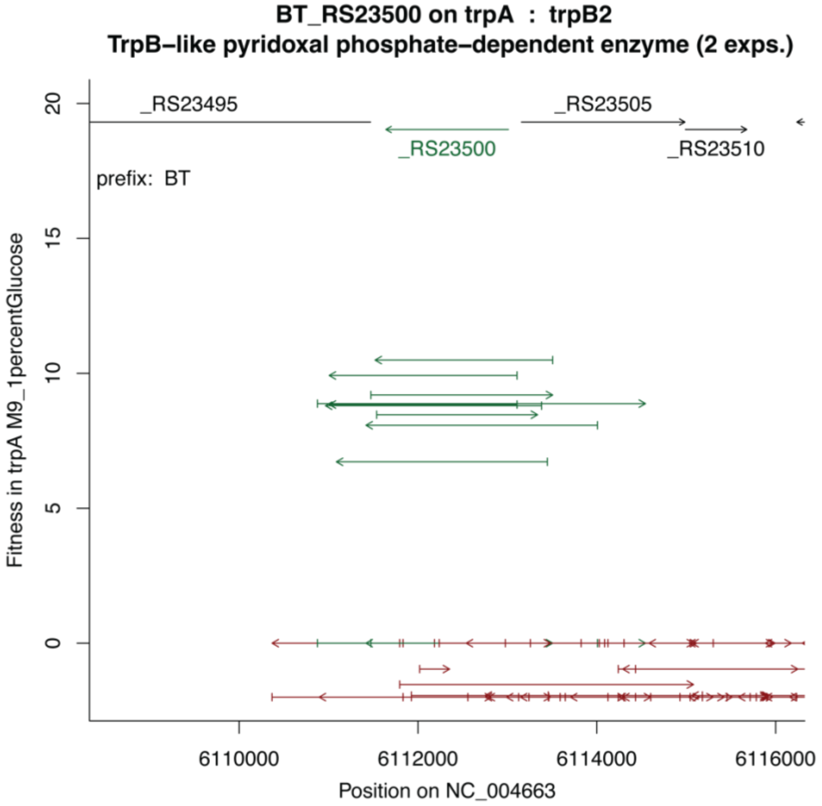


**Figure S29**. *Fitness benefit of BT_RS23500 of B. thetaiotaomicron associated fragments in the context of ΔtrpA.*

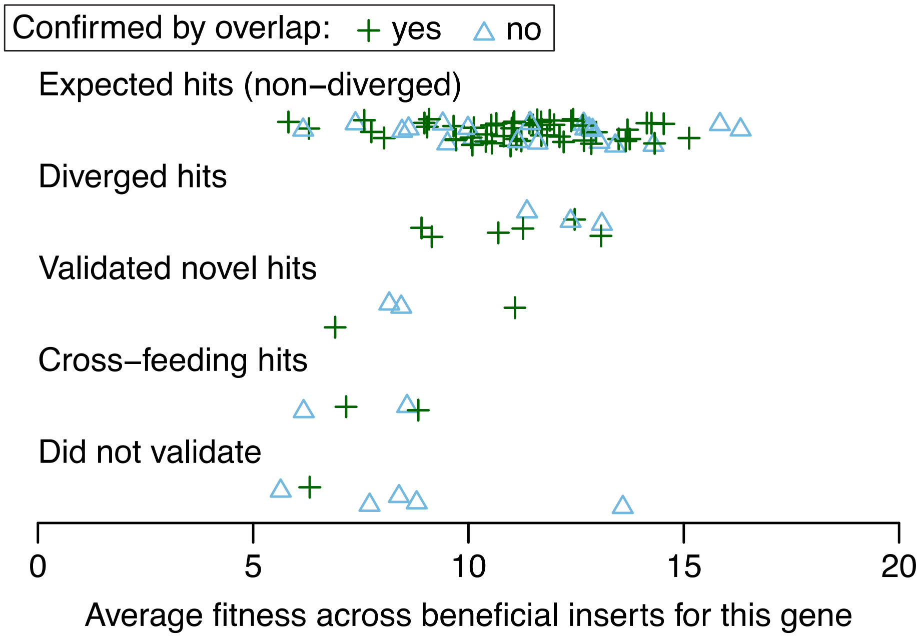


**Figure S30**. *Average fitness and overlap number for different categories of hits and outcomes of testing.* The trend that can be generally observed is that the hits that were not validated are more likely to have lower fitness values (closer to 5) and few to no overlapping fragments. For those that did not validate only one had a single overlapping fragment.

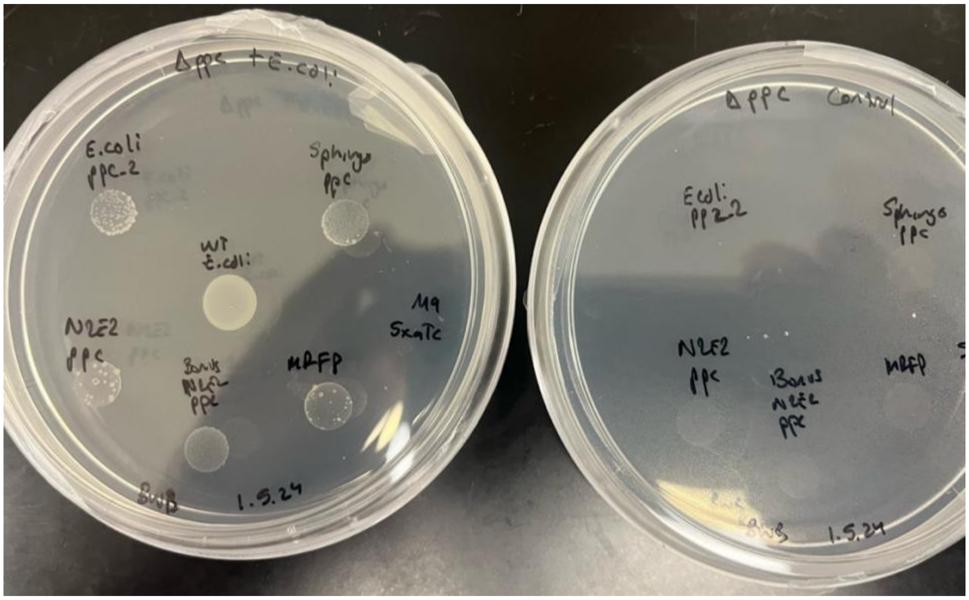


**Figure S31.** *Cross feeding with Δppc fitness enhancing fragments (dicarboxylic acid transporters)*. In the image are spotting assays of high-confidence hits (dicarboxylic acid transporters) identified in the context of Δ*ppc* rescue, spotted with (plate on the left) and without (plate on the right) wild-type *E. coli* on the same plate (spotted in the center). Candidate enzymes include BW25113_RS18290 of *E. Coli* ("E. Coli ppc_2", 10 o'clock), BDW16_RS00990 of *S. koreensis* ("Sphingo ppc", 2 o'clock), TK06_RS28425 of *P. fluorescens* FW300-N2E2 ("N2E2 ppc", 8 o'clock), along with a repeat of this condition that had better transformation efficiency ("Bonus N2E2 ppc", 6 o'clock). The control ("mRFP) can be found at 4 o'clock. As can be seen, there is not only a leading edge towards the wild-type *E. coli*, but the control (mRFP) lacks similar growth. This suggests that the wild-type *E. coli* is secreting the required nutrient to allow for grow of this knockout strain. And while it is possible for the control to take up this nutrient (see some colonies on the mRFP control), there is a significant benefit to overexpression of the transporters (additional growth, leading edge). None of the strains grow without the wild-type *E. coli* spotted (plate on the right).

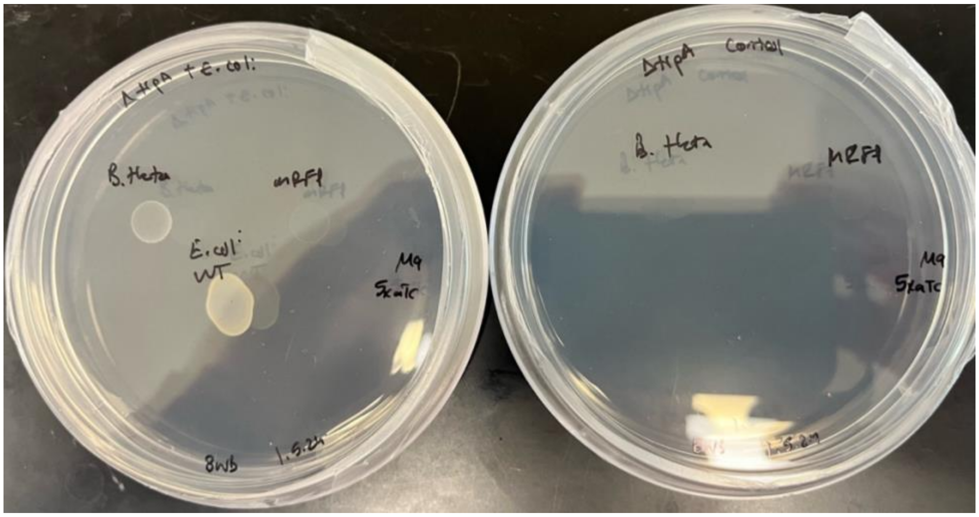


**Figure S32.** *Cross feeding in the context of the ΔtrpA fitness enhancing fragment*. Similar to the ppc high-confidence candidates, it can be seen that spotting the trpA candidate with (left) and without (right) wild-type *E. coli* shows some level cross feeding that allows for rescued growth. For the plate on the left, the strain containing the trpB2 candidate BT_RS23500 of *B. thetaiotaomicron* ("B. theta", 10 o'clock) shows growth while the control ("mRFP", 2 o'clock) does not. Without *E. coli* spotted, plate on the right, neither strain grows. As the proposed function of the trpB2 gene is use of indole to synthesize tryptophan, it is likely that indole is being secreted by wild-type *E. coli* and taken up by the strain overexpressing the trpB2 candidate from *B. thetaiotaomicron*, as direct provision of tryptophan itself would likely allow for the control to grow as well.

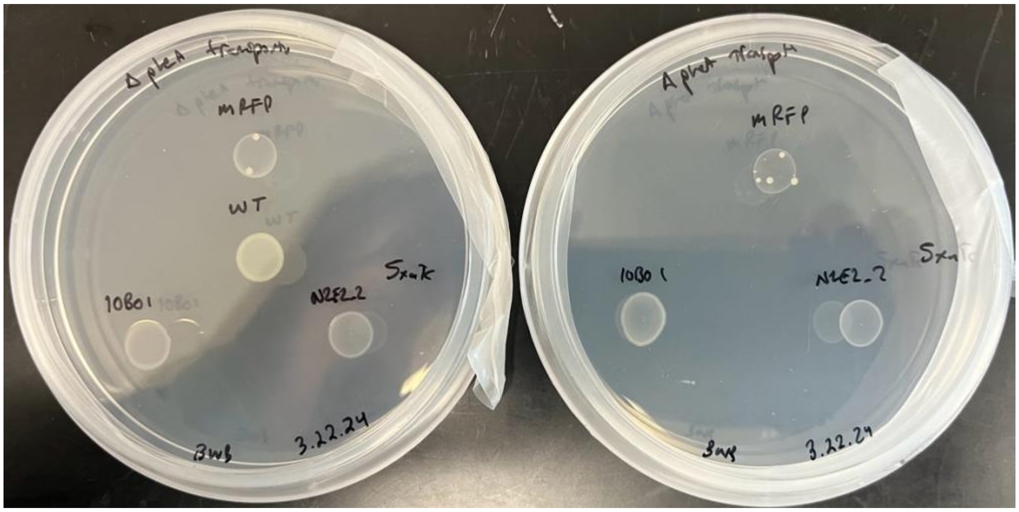


**Figure S33.** *Cross feeding in the context of ΔpheA fitness enhancing fragments containing putative MFS transporters LRK54_RS17455 and TK06_RS20405*. Spotting assays with LRK54_RS17455 of *Rhodanobacter denitrificans* FW104-10B01, TK06_RS20405 of *P. fluorescens* FW300-N2E2, and the mRFP control both with and without wild-type *E. coli* also spotted on the plate. Through the overall growth seems higher for the two MFS transporter candidate overexpressing strains compared to the mRFP control, the pattern is not differentiable between the cases where *E. coli* has and has not been co-spotted. Thus, it is not possible to confirmed a benefit of cross feeding.

### Supplemental Tables

**Table S1**. *Summary Table of genome fragment libraries generated. Cfus column represents colony forming units based on plating after transformation on LB chloramphenicol agar plates (selective for plasmid transformation). "BarSeq" represents the number of unique barcodes as identified by BarSeq sequencing. "PB Read #" is the total number of reads for that library during PacBio long-read sequencing. "PB Size" is the average size of the insert as identified by PacBio sequencing. "Gene #" is the total number of genes on the reference genome. "Gene Cov." Is the percentage of genes on the genome covered by the generated fragment library.*

| Strain | cfus | BarSeq | PB Read # | PB Size | Gene # | Gene Cov. |
|---|---|---|---|---|---|---|
| *Escherichia coli* (BW25113) | 66,373 | 56,500 | 954,803 | 2,628 | 4,233 | 97.54% |
| *Sphingomonas koreensis* (JSS26, DSMZ 15582) | 98,294 | 34,200 | 352,158 | 2,424 | 4,167 | 91.41% |
| *Bacillus subtilis* (168) | 82,333 | 68,700 | 506,000 | 2,300 | 4,240 | 90.35% |
| *Bacteroides thetaiotaomicron* (VPI-5482) | 55,520 | 52,000 | 497,338 | 2,074 | 4,682 | 54.57% |
| *Pseudomonas fluorescens* (FW300-N2E2) | 44,700 | 39,700 | 476,967 | 2,695 | 6,014 | 96.56% |
| *Lysobacter sp.* (FW306-1B-D06B) | 52,000 | 18,900 | 85,984 | 1,976 | 3,818 | 64.67% |
| *Xylophilus sp.* (GW821-FHT01B05) | 58,500 | 13,500 | 435,755 | 1,981 | 5,311 | 51.16% |
| *Rhodanobacter denitrificans* (FW104-10B01) | 84,500 | 46,200 | 355,863 | 2,108 | 3,534 | 82.00% |
| *Rhodoferax sp.* (GW822-FHT02A01) | 79,950 | 20,900 | 569,040 | 2,075 | 5,121 | 80.65% |
| *Pedobacter sp.* (FW305-3-2-15-E-R2A2) | 65,000 | 31,200 | 550,499 | 2,257 | 6,256 | 42.50% |
| *Acidovorax sp.* (FHTAMBA) | 97,500 | 52,700 | 336,044 | 2,192 | 4,344 | 90.24% |

**Table S2.** *Table of E. coli knockout auxotrophic strain backgrounds and missing function.*

| Knockout | Pathway | Function |
|---|---|---|
| ΔaroA | chorismate biosynthesis (amino acid) | 3-phosphoshikimate 1-carboxyvinyltransferase |
| ΔthrB | L-threonine biosynthesis (amino acid) | Homoserine kinase |
| ΔmetB | L-methionine biosynthesis (amnio acid) | Cystathionine gamma-synthase |
| ΔproB | L-proline biosynthesis (amino acid) | Glutamate 5-kinase |
| ΔproA | L-proline biosynthesis (amino acid) | Gamma-glutamyl phosphate reductase |
| ΔcysA | sulfate transport (aminio acid) | Sulfate/thiosulfate import ATP-binding protein CysA |
| ΔargG | L-arginine biosynthesis (amino acid) | Argininosuccinate synthase |
| ΔleuA | L-leucine biosynthesis (amino acid) | 2-isopropylmalate synthase |
| ΔtrpA | L-tryptophan biosynthesis (amino acid) | Tryptophan synthase alpha chain |
| ΔserA | L-serine biosynthesis (amino acid) | D-3-phosphoglycerate dehydrogenase |
| ΔhisC | L-histidine biosynthesis (amino acid) | Histidinol-phosphate aminotransferase |
| ΔmetE | L-methionine biosynthesis (amnio acid) | 5-methyltetrahydropteroyltriglutamate--homocysteine methyltransferase |
| ΔilvD | valine/isoleucine biosynthesis (amino acid) | Dihydroxy-acid dehydratase |
| ΔpheA | tyrosine/phenylalanine biosynthesis (amino acid) | Bifunctional chorismate mutase/prephenate dehydratase |
| ΔpurE | inosine-5'-phosphate biosynthesis (nucleotide) | N5-carboxyaminoimidazole ribonucleotide mutase |
| ΔcysH | assimilatory sulfate reduction (amino acid) | Phosphoadenosine 5'-phosphosulfate reductase |
| ΔhisG | L-histidine biosynthesis (amino acid) | ATP phosphoribosyltransferase |
| ΔaroE | chorismate biosynthesis (amino acid) | Shikimate dehydrogenase (NADP(+)) |
| ΔpyrD | UMP biosynthesis (nucleotide) | Dihydroorotate dehydrogenase (quinone) |
| Δppc | Mixed acid fermentation | Phosphoenolpyruvate carboxylase |

**Table S3.** *Summary of all transformations run.*

| Name | Fragment Library | KO | Transformation Efficiency (cfus) |
|---|---|---|---|
| BBL1 | pBWBHTC5, *Bacteroides thetaiotaomicron* | ΔthrB | 170,000 |
| BBL2 | pBWBHTC5, *Bacteroides thetaiotaomicron* | ΔaroA | 73,000 |
| BBL3 | pBWBHTC4, *Bacillus subtilis* 168 | ΔthrB | 178,000 |
| BBL4 | pBWBHTC4, *Bacillus subtilis* 168 | ΔaroA | 80,000 |
| BBL5 | pBWBHTC5, *Bacteroides thetaiotaomicron* | ΔmetB | 9750000 |
| BBL6 | pBWBHTC5, *Bacteroides thetaiotaomicron* | ΔproA | 14300000 |
| BBL7 | pBWBHTC4, *Bacillus subtilis* 168 | ΔmetB | 40300000 |
| BBL8 | pBWBHTC4, *Bacillus subtilis* 168 | ΔproA | 7150000 |
| BBL9 | pBWBHTC1, *Escherichia coli* BW25113 | ΔmetB | 1592500 |
| BBL10 | pBWBHTC1, *Escherichia coli* BW25113 | ΔproA | 5200000 |

| | | | |
|---|---|---|---|
| BBL11 | pBWBHTC2, *Pseudomonas fluorescens* (FW300-N2E2) | ΔmetB | 1560000 |
| BBL12 | pBWBHTC2, *Pseudomonas fluorescens* (FW300-N2E2) | ΔproA | 975000 |
| BBL13 | pBWBHTC3, *Sphingomonas koreensis* | ΔproA | 3900000 |
| BBL14 | pBWBHTC6, BWB514, FW306-1B-D06B, *Lysobacter sp*. | ΔproA | 5200000 |
| BBL15 | pBWBHTC7, BWB549, GW821-FHT01B05, *Xylophilus sp*. | ΔproA | 3445000 |
| BBL16 | pBWBHTC8, BWB510, FW104-10B01, *Rhodanobacter denitrificans* | ΔproA | 3683333 |
| BBL17 | pBWBHTC9 - BWB553 GW822-FHT02A01, *Rhodoferax sp*. | ΔproA | 26000 |
| BBL18 | pBWBHTC10 - BWB503, FW305-3-2-15-E-R2A2, *Pedobacter sp*. | ΔproA | 1300000 |
| BBL19 | pBWBHTC11 - BWB602, FHTAMBA, *Acidovorax sp*. | ΔproA | 6500 |
| BBL20 | pBWBHTC3 - *Sphingomonas koreensis* | ΔmetB | 17550000 |
| BBL21 | pBWBHTC6, BWB514, FW306-1B-D06B, *Lysobacter sp*. | ΔmetB | 5200000 |
| BBL22 | pBWBHTC7, BWB549, GW821-FHT01B05, *Xylophilus sp*. | ΔmetB | 2600000 |
| BBL23 | pBWBHTC8, BWB510, FW104-10B01, *Rhodanobacter denitrificans* | ΔmetB | 5850000 |
| BBL24 | pBWBHTC9 - BWB553 GW822-FHT02A01, *Rhodoferax sp*. | ΔmetB | 130000 |
| BBL25 | pBWBHTC10 - BWB503, FW305-3-2-15-E-R2A2, *Pedobacter sp*. | ΔmetB | 19500000 |
| BBL26 | pBWBHTC11 - BWB602, FHTAMBA, *Acidovorax sp*. | ΔmetB | 4875000 |
| BBL27 | pBWBHTC1, *Escherichia coli* BW25113 | ΔaroA | 1170000 |
| BBL28 | pBWBHTC2, *Pseudomonas fluorescens* (FW300-N2E2) | ΔaroA | 4268333.333 |
| BBL29 | pBWBHTC3, *Sphingomonas koreensis* | ΔaroA | 17225000 |
| BBL30 | pBWBHTC6, BWB514, FW306-1B-D06B, *Lysobacter sp*. | ΔaroA | 2600000 |
| BBL31 | pBWBHTC7, BWB549, GW821-FHT01B05, *Xylophilus sp*. | ΔaroA | 10075000 |
| BBL32 | pBWBHTC8, BWB510, FW104-10B01, *Rhodanobacter denitrificans* | ΔaroA | 3250000 |
| BBL33 | pBWBHTC9 - BWB553 GW822-FHT02A01, *Rhodoferax sp*. | ΔaroA | 39000 |
| BBL34 | pBWBHTC10 - BWB503, FW305-3-2-15-E-R2A2, *Pedobacter sp*. | ΔaroA | 17550000 |

| | | | |
|---|---|---|---|
| BBL35 | pBWBHTC11 - BWB602, FHTAMBA, *Acidovorax sp.* | ΔaroA | 1495000 |
| BBL36 | pBWBHTC1, *Escherichia coli* BW25113 | ΔthrB | 585000 |
| BBL37 | pBWBHTC2, *Pseudomonas fluorescens* (FW300-N2E2) | ΔthrB | 910000 |
| BBL38 | pBWBHTC3, *Sphingomonas koreensis* | ΔthrB | 1657500 |
| BBL39 | pBWBHTC6, BWB514, FW306-1B-D06B, *Lysobacter sp.* | ΔthrB | 910000 |
| BBL40 | pBWBHTC7, BWB549, GW821-FHT01B05, *Xylophilus sp.* | ΔthrB | 1137500 |
| BBL41 | pBWBHTC8, BWB510, FW104-10B01, *Rhodanobacter denitrificans* | ΔthrB | 1657500 |
| BBL42 | pBWBHTC9 - BWB553 GW822-FHT02A01, *Rhodoferax sp.* | ΔthrB | 19500 |
| BBL43 | pBWBHTC10 - BWB503, FW305-3-2-15-E-R2A2, *Pedobacter sp.* | ΔthrB | 6500000 |
| BBL44 | pBWBHTC11 - BWB602, FHTAMBA, *Acidovorax sp.* | ΔthrB | 2632500 |
| BBL45 | Mix of all 11 | ΔthrB | 910000 |
| BBL46 | Mix of all 11 | ΔaroA | 81250 |
| BBL47 | Mix of all 11 | ΔmetB | 1235000 |
| BBL48 | Mix of all 11 | ΔproA | 780000 |
| BBL49 | Mix of all 11 | ΔCysA | 113750 |
| BBL50 | Mix of all 11 | ΔArgG | 6175000 |
| BBL51 | Mix of all 11 | ΔLeuA | 487500 |
| BBL52 | Mix of all 11 | ΔtrpA | 877500 |
| BBL53 | Mix of all 11 | ΔserA | 975000 |
| BBL54 | Mix of all 11 | ΔhisC | 5200000 |
| BBL55 | Mix of all 11 | ΔpheA | 16900000 |
| BBL56 | Mix of all 11 | ΔpurE | 2827500 |
| BBL57 | Mix of all 11 | Δppc | 1170000 |
| BBL58 | Mix of all 11 | ΔproB | 650000 |
| BBL59 | Mix of all 11 | ΔcysH | 325000 |
| BBL60 | Mix of all 11 | ΔmetE | 5525000 |
| BBL61 | Mix of all 11 | ΔserB | 2925000 |
| BBL62 | Mix of all 11 | ΔhisG | 1040000 |
| BBL63 | Mix of all 11 | ΔaroE | 2210000 |
| BBL64 | Mix of all 11 | ΔpyrD | 1072500 |
| BBL65 | Mix of all 11 | ΔilvD | 1690000 |

**Table S4.** *Summary of all selections.*

| Name | Fragment Library | Reference Transformation | KO destination | aTc concentration |
|---|---|---|---|---|
| BBS1 | pBWBHTC5 - *Bacteroides thetaiotaomicron* | BBL2 | ΔaroA | 5x |
| BBS2 | pBWBHTC4, *Bacillus subtilis* 168 | BBL3 | ΔthrB | 5x |
| BBS3 | pBWBHTC4, *Bacillus subtilis* 168 | BBL4 | ΔaroA | 5x |
| BBS4 | pBWBHTC5 - *Bacteroides thetaiotaomicron* | BBL2 | ΔaroA | 0.5x |
| BBS5 | pBWBHTC5 - *Bacteroides thetaiotaomicron* | BBL2 | ΔaroA | 10x |
| BBS6 | pBWBHTC4, *Bacillus subtilis* 168 | BBL3 | ΔthrB | 0.5x |
| BBS7 | pBWBHTC4, *Bacillus subtilis* 168 | BBL3 | ΔthrB | 10x |
| BBS8 | pBWBHTC4, *Bacillus subtilis* 168 | BBL4 | ΔaroA | 0.5x |
| BBS9 | pBWBHTC4, *Bacillus subtilis* 168 | BBL4 | ΔaroA | 10x |
| BBS10 | pBWBHTC4, *Bacillus subtilis* 168 | BBL7 | ΔmetB | 1x |
| BBS11 | pBWBHTC4, *Bacillus subtilis* 168 | BBL7 | ΔmetB | 5x |
| BBS12 | pBWBHTC4, *Bacillus subtilis* 168 | BBL7 | ΔmetB | 10x |
| BBS13 | pBWBHTC4, *Bacillus subtilis* 168 | BBL8 | ΔproA | 1x |
| BBS14 | pBWBHTC4, *Bacillus subtilis* 168 | BBL8 | ΔproA | 10x |
| BBS15 | pBWBHTC1, *Escherichia coli* BW25113 | BBL10 | ΔproA | 1x |
| BBS16 | pBWBHTC1, *Escherichia coli* BW25113 | BBL9 | ΔmetB | 1x |
| BBS17 | pBWBHTC1, *Escherichia coli* BW25113 | BBL9 | ΔmetB | 5x |
| BBS18 | pBWBHTC8, BWB510, FW104-10B01, *Rhodanobacter denitrificans* | BBL16 | ΔproA | 1x |
| BBS19 | pBWBHTC8, BWB510, FW104-10B01, *Rhodanobacter denitrificans* | BBL16 | ΔproA | 5x |
| BBS20 | pBWBHTC7, BWB549, GW821-FHT01B05, *Xylophilus sp*. | BBL15 | ΔproA | 1x, 5x |
| BBS21 | pBWBHTC2, *Pseudomonas fluorescens* (FW300-N2E2) | BBL12 | ΔproA | 1x, 5x |
| BBS22 | pBWBHTC6 - BWB514, FW306-1B-D06B, *Lysobacter sp*. | BBL14 | ΔproA | 1x, 5x |
| BBS23 | pBWBHTC5 - *Bacteroides thetaiotaomicron* | BBL5 | ΔmetB | 1x |
| BBS24 | pBWBHTC5 - *Bacteroides thetaiotaomicron* | BBL6 | ΔproA | 1x, 5x, 10x |
| BBS25 | pBWBHTC3, *Sphingomonas koreensis* | BBL13 | ΔproA | 5x |
| BBS26 | pBWBHTC11 - BWB602, FHTAMBA, *Acidovorax sp*. | BBL26 | ΔmetB | 1x, 5x |

| | | | | |
|---|---|---|---|---|
| BBS27 | pBWBHTC9 - BWB553 GW822-FHT02A01, *Rhodoferax sp*. | BBL24 | ΔmetB | 1x, 5x |
| BBS28 | pBWBHTC10 - BWB503, FW305-3-2-15-E-R2A2, *Pedobacter sp*. | BBL18 | ΔproA | 1x, 5x |
| BBS29 | pBWBHTC11 - BWB602, FHTAMBA, *Acidovorax sp*. | BBL19 | ΔproA | 1x, 5x |
| BBS30 | pBWBHTC9 - BWB553 GW822-FHT02A01, *Rhodoferax sp*. | BBL17 | ΔproA | 1x, 5x |
| BBS31 | pBWBHTC6 - BWB514, FW306-1B-D06B, *Lysobacter sp*. | BBL21 | ΔmetB | 1x, 5x |
| BBS32 | pBWBHTC1, *Escherichia coli* BW25113 | BBL27 | ΔaroA | 1x, 5x |
| BBS33 | pBWBHTC2, *Pseudomonas fluorescens* (FW300-N2E2) | BBL28 | ΔaroA | 1x, 5x |
| BBS34 | pBWBHTC8, BWB510, FW104-10B01, *Rhodanobacter denitrificans* | BBL32 | ΔaroA | 1x, 5x |
| BBS35 | pBWBHTC10 - BWB503, FW305-3-2-15-E-R2A2, *Pedobacter sp*. | BBL34 | ΔaroA | 1x, 5x |
| BBS36 | pBWBHTC11 - BWB602, FHTAMBA, *Acidovorax sp*. | BBL35 | ΔaroA | 1x, 5x |
| BBS37 | pBWBHTC8, BWB510, FW104-10B01, *Rhodanobacter denitrificans* | BBL23 | ΔmetB | 5x |
| BBS38 | pBWBHTC1, Escherichia coli BW25113 | BBL36 | ΔthrB | 5x |
| BBS39 | pBWBHTC2, *Pseudomonas fluorescens* (FW300-N2E2) | BBL37 | ΔthrB | 5x |
| BBS40 | pBWBHTC6 - BWB514, FW306-1B-D06B, *Lysobacter sp*. | BBL39 | ΔthrB | 1x, 5x |
| BBS41 | pBWBHTC7, BWB549, GW821-FHT01B05, *Xylophilus sp*. | BBL40 | ΔthrB | 5x |
| BBS42 | pBWBHTC3, *Sphingomonas koreensis* | BBL38 | ΔthrB | 1x, 5x |
| BBS43 | Mix -11 | BBL45 | ΔthrB | 1x |
| BBS44 | Mix -11 | BBL45 | ΔthrB | 5x |
| BBS45 | Mix -11 | BBL47 | ΔmetB | 1x |
| BBS46 | Mix -11 | BBL47 | ΔmetB | 5x |
| BBS47 | pBWBHTC9 - BWB553 GW822-FHT02A01, *Rhodoferax sp*. | BBL33 | ΔaroA | 5x |
| BBS48 | pBWBHTC8, BWB510, FW104-10B01, *Rhodanobacter denitrificans* | BBL41 | ΔthrB | 1x, 5x |
| BBS49 | Mix - 11 | BBL48 | ΔproA | 1x, 5x |
| BBS50 | Mix - 11 | BBL46 | ΔaroA | 1x |
| BBS51 | pBWBHTC11 - BWB602, FHTAMBA, *Acidovorax sp*. | BBL44 | ΔthrB | 1x, 5x |

| | | | | |
|---|---|---|---|---|
| BBS52 | pBWBHTC9 - BWB553 GW822-FHT02A01, *Rhodoferax sp.* | BBL42 | ΔthrB | 1x, 5x |
| BBS53 | Mix - 11 | BBL46 | ΔaroA | 5x |
| BBS54 | Mix - 11 | BBL49 | ΔCysA | 1x |
| BBS55 | Mix - 11 | BBL49 | ΔCysA | 5x |
| BBS56 | Mix - 11 | BBL51 | ΔleuA | 1x |
| BBS57 | Mix - 11 | BBL51 | ΔleuA | 5x |
| BBS58 | Mix - 11 | BBL52 | ΔtrpA | 1x |
| BBS59 | Mix - 11 | BBL52 | ΔtrpA | 5x |
| BBS60 | Mix - 11 | BBL50 | ΔargA | 1x |
| BBS61 | Mix - 11 | BBL50 | ΔargA | 5x |
| BBS62 | Mix - 11 | BBL53 | ΔserA | 1x |
| BBS63 | Mix - 11 | BBL53 | ΔserA | 5x |
| BBS64 | Mix - 11 | BBL55 | ΔpheC | 1x |
| BBS65 | Mix - 11 | BBL55 | ΔpheC | 5x |
| BBS66 | Mix - 11 | BBL56 | ΔpurE | 1x |
| BBS67 | Mix - 11 | BBL56 | ΔpurE | 5x |
| BBS68 | Mix - 11 | BBL54 | ΔhisC | 1x |
| BBS69 | Mix - 11 | BBL54 | ΔhisC | 5x |
| BBS70 | Mix - 11 | BBL57 | Δppc | 1x |
| BBS71 | Mix - 11 | BBL57 | Δppc | 5x |
| BBS72 | Mix - 11 | BBL58 | ΔproB | 1x |
| BBS73 | Mix - 11 | BBL58 | ΔproB | 5x |
| BBS74 | Mix - 11 | BBL60 | ΔmetE | 1x |
| BBS75 | Mix - 11 | BBL60 | ΔmetE | 5x |
| BBS76 | Mix - 11 | BBL61 | ΔserB | 1x |
| BBS77 | Mix - 11 | BBL61 | ΔserB | 5x |
| BBS78 | Mix - 11 | BBL63 | ΔaroE | 1x |
| BBS79 | Mix - 11 | BBL63 | ΔaroE | 5x |
| BBS80 | Mix - 11 | BBL64 | ΔpyrD | 1x |
| BBS81 | Mix - 11 | BBL64 | ΔpyrD | 5x |
| BBS82 | Mix - 11 | BBL65 | ΔilvD | 1x |
| BBS83 | Mix - 11 | BBL65 | ΔilvD | 5x |
| BBS84 | Mix - 11 | BBL62 | ΔhisG | 1x |
| BBS85 | Mix - 11 | BBL62 | ΔhisG | 5x |
| BBS86 | Mix - 11 | BBL59 | ΔcysH | 1x |
| BBS87 | Mix - 11 | BBL59 | ΔcysH | 5x |

**Table S5**. *First of three tests for liquid culture selection. Escherichia coli into ΔthrB.*

| ***Escherichia coli* BW25113 into Δ*thrB*** | | |
|---|---|---|
| Barcode | Reads | Fraction of Total |
| CACTAGTTGAAGTGTGTGGG | 4359217 | 0.998328187 |
| TTGCAAAATAAATCCTTATT | 3396 | 0.000777737 |
| GCATGTTGCGGTGAAATAAG | 3191 | 0.000730788 |
| GCGCCTACTTACGGCTATTG | 419 | 9.59575E-05 |
| TTATTTTCGTAAGTTGTGCG | 122 | 2.79399E-05 |
| GTATGATCAGCGTCGCTGGG | 84 | 1.92373E-05 |
| GTGCAGGGCAGAGACATTGT | 45 | 1.03057E-05 |
| TTCTGGTCTCGTGGGAGCAC | 33 | 7.55751E-06 |
| GTTAGCTCGGGACAGACAAA | 10 | 2.29015E-06 |

**Table S6.** *Second of three tests of liquid culture outgrowth. Sphingomonas koreensis into ΔthrB.*

| ***Sphingomonas koreensis* into ΔthrB** | | |
|---|---|---|
| Barcode | Reads | Fraction of Total |
| TTCTGGTCTCGTGGGAGCAC | 4504153 | 0.977960804 |
| GCAACTTGTAGTCAGGAAAG | 101349 | 0.022005325 |
| TTATTTTCGTAAGTTGTGCG | 103 | 2.23638E-05 |
| CACTAGTTGAAGTGTGTGGG | 26 | 5.64523E-06 |
| GTTAGCTCGGGACAGACAAA | 11 | 2.38837E-06 |
| AAATAGTTTTGAGGGGTCCG | 4 | 8.68497E-07 |
| GCGGGGCTAGTCTAAAGCAG | 4 | 8.68497E-07 |
| GGCAAGTGTGTTATGCGTGG | 4 | 8.68497E-07 |
| GTCAGTTGAGCCTCGCTGCG | 4 | 8.68497E-07 |

**Table S7.** *Third of three tests of liquid culture outgrowth. Pseudomonas fluorescens into ΔthrB.*

| ***Pseudomonas fluorescens* FW300-N2E2 into ΔthrB** | | |
|---|---|---|
| Barcode | Reads | Fraction of Total |
| TTATTTTCGTAAGTTGTGCG | 4004179 | 0.846429397 |
| GTTAGCTCGGGACAGACAAA | 725435 | 0.153347168 |
| GTGTTCTTGTAACTCTGATT | 969 | 0.000204834 |
| CACTAGTTGAAGTGTGTGGG | 41 | 8.66685E-06 |
| TTCTGGTCTCGTGGGAGCAC | 35 | 7.39853E-06 |
| TATGCAACGGTATTGCTACT | 6 | 1.26832E-06 |
| AACAGGTGCGAGGCGTCTTT | 2 | 4.22773E-07 |
| GATCACCGACTTTTATACAC | 2 | 4.22773E-07 |
| GATTTTAAATTTGCCGGCGA | 2 | 4.22773E-07 |

*Table S8. Diverged hits.*

| Background | Source | locus_tag | protein_id | uniprotId | Expected | Comments |
|---|---|---|---|---|---|---|
| aroA | *Bacteroides thetaiotaomicron* VPI-5482 | BT_RS11065 | WP_162303115.1 | Q8A5Q2 | yes | 35% identical to the *Vibrio cholerae* enzyme (Q9KRB0) |
| aroE | *Sphingomonas koreensis* JSS26; DSMZ 15582 | BDW16_RS10815 | WP_066578130.1 | A0A2M8WD96 | yes | Also known as Ga0059261_2194; RB-TnSeq data confirms that mutants are auxotrophic |
| hisC | *Acidovorax* sp. FHTAMBA | AAFF19_05770 | XAH19868.1 | A0A2R7PAQ8 | yes | 32% identical to HisC from *Caldanaerobacter* (Q8R5Q4); considered expected because it is in an operon with histidine synthesis genes |
| hisC | *Rhodoferax sp.* GW822-FHT02A01 | AAGF34_01100 | XAF49772.1 | NA | no | 31% identical to HisC from *Caldanaerobacter* (Q8R5Q4); is in an operon with histidine synthesis genes |
| metB | *Bacillus subtilis* 168 | BSU_11870 | NP_389069.1 | A0A9Q4E7A0 | no | This is the *B. subtilis* O-acetylhomoserine sulfhydrylase (MetI); it was previously reported to complement metB- *E. coli*, even though *in vitro*, the enzyme has no detectable activity on O-succinylhomoserine (PMID:11832514) |
| proB | *Pedobacter sp*. FW305-3-2-15-E-R2A2 | AAFF35_11135 | WZV14794.1 | A0A1H0J514 | yes | 35% identical to CA265_RS20855, whose mutants are auxotrophic (RB-TnSeq data) |
| thrB | *Bacillus subtilis* 168 | BSU_32240 | NP_391104.1 | A0AA96UM70 | yes | Complementation was previously demonstrated (PMC1167255) but this information is not in the curated databases |
| trpA | *Pedobacter sp*. FW305-3-2-15-E-R2A2 | AAFF35_06155 | WZV13815.1 | NA | yes | 38% identical to the budding yeast enzyme (P00931) |
| trpA | *Acidovorax sp*. FHTAMBA | AAFF19_07375 | XAH20166.1 | H0BV16 | yes | 66% identical to RS_RS09955, whose mutant is auxotrophic for tryptophan (PMC8510521); this information is not in the curated databases |
| trpA | *Rhodanobacter denitrificans* FW104-10B01 | LRK54_RS01680 | WP_027489804.1 | M4NLA4 | yes | 39.7% identical to the maize enzyme (B2Y0K4) |

**Table S9.** *Follow Up tests.*

| KO | locus_tag | protDesc | curated | protein_id | uniprotId | Genome | t0set | Validated |
|---|---|---|---|---|---|---|---|---|
| cysA | BSU_15580 | sulfate permease | cysA* | NP_389441.1 | A0A6M4JJU8 | *Bacillus subtilis* (168) | Mix_11.cysA | Yes |
| cysA | TK06_RS10770 | sulfite exporter TauE/SafE family protein | cysA* | WP_063322060.1 | A0A1K1T875 | *Pseudomonas fluorescens* (FW300-N2E2) | Mix_11.cysA | Yes |
| hisC | TK06_RS12685 | histidinol-phosphate transaminase | NA | WP_063322357.1 | A0A159ZZQ5 | *Pseudomonas fluorescens* (FW300-N2E2) | Mix_11.hisC | Yes |
| metB | LRK54_RS05660 | PLP-dependent aspartate aminotransferase family protein | NA | WP_027489953.1 | A0A368KK22 | FW104-10B01, *Rhodanobacter denitrificans* | pBWBHTC8.metB | Yes |
| ppc | BW25113_RS18290 | C4-dicarboxylate transporter DctC | NA | WP_000858214.1 | D3H0Y2 | *Escherichia coli* BW25113 | Mix_11.ppc | Crossfed |
| ppc | BDW16_RS00990 | dicarboxylate/amino acid:cation symporter | NA | WP_066575250.1 | A0A1L6JC14 | *Sphingomonas koreensis* | Mix_11.ppc | Crossfed |
| ppc | TK06_RS28425 | dicarboxylate/amino acid:cation symporter | NA | WP_063324741.1 | A0A160A458 | *Pseudomonas fluorescens* (FW300-N2E2) | Mix_11.ppc | Crossfed |
| trpA | BT_RS23500 | TrpB-like pyridoxal phosphate-dependent enzyme | trpB2 | WP_008760379.1 | D7IA57 | *Bacteroides thetaiotaomicron* | Mix_11.trpA | Crossfed |
| pheA | LRK54_RS17455 | MFS transporter | NA | WP_063090425.1 | M4NEK0 | FW104-10B01, *Rhodanobacter denitrificans* | Mix_11.pheA | No |
| pheA | TK06_RS20405 | MFS transporter | NA | WP_063323551.1 | A0A160A0V0 | *Pseudomonas fluorescens* (FW300-N2E2) | Mix_11.pheA | No |
| proB | BW25113_RS20525 | phosphoenolpyruvate carboxylase | ppc | WP_001005586.1 | C5A0C2 | *Escherichia coli* BW25113 | Mix_11.ppc | No |
| proB | TK06_RS22845 | dihydroxy-acid dehydratase | ilvD | WP_063323942.1 | A0A160A1W0 | *Pseudomonas fluorescens* (FW300-N2E2) | Mix_11.proB | No |
| pyrD | TK06_RS26365 | SDR family oxidoreductase | NA | WP_063324432.1 | A0A165ZNW9 | *Pseudomonas fluorescens* (FW300-N2E2) | Mix_11.pyrD | No |
| cysH | BSU_32380 | putative transporter | NA | NP_391118.1 | A0A6H0H720 | *Bacillus subtilis* (168) | Mix_11.cysH | No |
